# ATP Citrate Lyase Drives Vascular Remodeling Diseases Development Through Metabolic-Epigenetic Reprograming

**DOI:** 10.1101/2024.02.02.578545

**Authors:** Yann Grobs, Charlotte Romanet, Sarah-Eve Lemay, Alice Bourgeois, Pierre Voisine, Charlie Theberge, Melanie Sauvaget, Sandra Breuils-Bonnet, Sandra Martineau, Reem El Kabbout, Manon Mougin, Elizabeth Dumais, Jean Perron, Nicolas Flamand, François Potus, Steeve Provencher, Olivier Boucherat, Sebastien Bonnet

**Author notes:** Corresponding authors: Sébastien Bonnet, Ph.D. Pulmonary Hypertension Research Group, Québec Heart and Lung Institute Research Centre 2725, chemin Sainte-Foy, Québec, QC Canada, G1V 4G5. These authors contributed equally to this work (Yann Grobs, Charlotte Romanet). These authors co-supervised the study (Olivier Boucherat, Sebastien Bonnet).

## Abstract

Our study explores the previously uncharted role of ATP-citrate lyase (ACLY) in vascular remodeling within the pulmonary and coronary arteries, providing novel insights into the pathogenesis of pulmonary hypertension and coronary artery diseases. ACLY, involved in de novo lipid synthesis and histone acetylation, has emerged as a key regulator in sustaining vascular smooth muscle cell (VSMC) proliferation and survival.

Utilizing human coronary and pulmonary artery tissues, our findings reveal an upregulation of ACLY expression during vascular remodeling processes. Inhibition of ACLY, achieved through pharmacological and molecular interventions in humans primary cultured VSMCs, leads to decreased proliferation, migration, and resistance to apoptosis. Mechanistically, these effects are associated with diminished glycolysis, lipid synthesis, GCN5-dependent histone acetylation, and FOXM1 activation.

In vivo experiments, combining pharmacological and VSMC-specific ACLY knockout mice, ACLY inhibition demonstrates its efficacy in mitigating coronary artery remodeling and reducing pulmonary hypertension. Notably, initiating ACLY inhibition post-disease onset reverses pathological conditions, positioning ACLY as a promising therapeutic target.

Human ex vivo tissue culture further supports our findings, showing reduced vascular remodeling in cultured human coronary artery rings and a reversal of pulmonary artery remodeling in precision-cut lung slices upon ACLY inhibition. This study introduces a groundbreaking concept, linking disparate abnormalities in vascular diseases to a common pathogenetic denominator, ACLY. The identified “multiple hit” therapeutic approach presents potential targets for addressing complex vascular diseases, offering avenues for future clinical interventions.

**ONE SENTENCE SUMMARY:** Our study delineates the pivotal role of ATP-citrate lyase in orchestrating vascular remodeling, establishing it as a compelling translational target for therapeutic interventions in pulmonary hypertension and coronary artery disease.

## INTRODUCTION

Arterial remodeling describes the structural changes that occur within the vascular wall as a result of a wide range of physiological and pathophysiological triggers (*1*). While this remodeling is a natural aspect of aging, premature arterial remodeling is a hallmark of many cardiovascular and pulmonary diseases, including atherosclerosis, post-angioplasty restenosis, and pulmonary arterial hypertension (PAH). In these conditions, the proliferation of vascular smooth muscle cells (VSMCs) is abnormally high, which is a critical factor in the disease process.

Due to this shared pathogenic mechanism, there is a notable overlap in histopathological features among these distinct medical conditions. Consequently, this indicates that innovative treatments developed for one disorder have the potential to be repurposed effectively to manage the others. This insight into shared disease pathways opens up promising avenues for cross-applicable therapeutic strategies, enhancing the prospect of improving patient outcomes across a spectrum of cardiopulmonary diseases.

Despite considerable progress towards understanding the signaling pathways implicated in the structural alterations of the arterial wall and advances in medical therapy, cardiovascular diseases remain the leading cause of morbidity and mortality worldwide, stressing the urgent need for more effective therapeutic approaches. Moreover, given the complexity and multifactorial nature of pathological remodeling in cardiopulmonary disorders, it is of great therapeutic interest to identify and characterize new actionable targets that intersect with multiple detrimental cellular stress responses.

Vascular SMC phenotypic changes, resulting in excessive survival and proliferation, are prominent features of neointimal and medial lesions seen in systemic and pulmonary vascular diseases. It is now well-established that a dysregulated metabolism with the preferential use of aerobic glycolysis rather than oxidative phosphorylation coupled to epigenetic changes create a feedback loop that synergistically nurtures the abnormal phenotype of arterial cells (*2–5*). In view of this, factors critically involved in this abnormal metabolism-epigenetics crosstalk represent attractive targets for a therapeutic approach.

Acetyl-Coenzyme A (acetyl-CoA) occupies a critical position at the crossroads of metabolism and chromatin dynamics. Generated from many precursors such as citrate, pyruvate or acetate, cytosolic acetyl-CoA serves as a building block for fatty acids (FA) and cholesterol synthesis, while in the nucleus, it provides acetyl groups necessary for histone acetylation and thus gene expression regulation (*6*). By converting mitochondria-derived citrate into acetyl-CoA, the nucleo-cytoplasmic enzyme ATP citrate lyase (ACLY) plays a key role in the crosstalk between metabolism and epigenetic modifications. In addition to provide the cell with membrane lipids required for cell proliferation (*7*), compelling evidence has shown that ACLY is essential for sustaining aberrant survival and proliferation of cancer cells by multiple intertwined mechanisms(*8*). Indeed, nuclear production of acetyl-CoA by ACLY was documented to enhance site-specific histone acetylation and oncogenic gene expression programs by providing the substrate for histone acetyltransferases(*9*). Besides its impact on gene expression, activation of ACLY contributes to maintain the cytosolic gauge of citrate at a low level, which sustains the Warburg effect (*8*). In this regard, “turning off” ACLY has repeatedly shown to elicit anti-cancer effects in preclinical models (*10, 11*) and pharmacological ACLY inhibitors, previously developed and approved by the FDA as a lipid lowering drugs, have attracted a growing interest as promising anti-cancer agents(*10, 11*).

Given that vascular SMCs in vascular remodeling diseases exhibit behavior akin to cancer cells (*10, 11*), ACLY appears to be an exciting therapeutic target, inhibition of which may deliver a combined attack on multiple mechanisms implicated in pathological vascular remodeling. Surprisingly, its role in vascular SMC hyperplasia seen in both PAH and coronary artery disease (CAD), remains unknown. In the present study, we used human tissues and cells as well as a combination of genetic and pharmacological approaches in animal models to unveil a pivotal role of ACLY in these vascular remodeling diseases.

## RESULTS

### Increased expression of ACLY in human PAH and CAD

ACLY expression is regulated at multiple levels, with phosphorylation at serine 455 serving as a mechanism to enhance its activity(*12, 13*). As a starting point to study the role of ACLY in vascular remodeling diseases, we first measured the expression of its phosphorylated and total forms in dissected pulmonary arteries (PAs) and isolated PASMCs from control and PAH patients. Western blot analysis revealed a significant increase in both pACLY and ACLY expression in PAH tissues and PAH-PASMCs (Fig. 1, A and B). Correspondingly, ACLY immunoreactivity was increased in remodeled small PAs from PAH patients (Supplemental Fig. S1A). Consistent with findings in human PAH, the expression levels of pACLY and ACLY were notably heightened in CoASMCs isolated from patients with CAD compared to controls (Fig. 1C). Immunofluorescence analyses further supported this observation by demonstrating an upregulation of ACLY expression in remodeled coronary arteries (CoA) from CAD patients (Fig. S1B). While ACLY is acknowledged as the primary producer of acetyl-CoA, alternative enzymes like acetyl-CoA synthase 2 (ACSS2) have the capacity to generate acetyl-CoA from acetate. We assessed ACSS2 expression in both PAH-PASMCs and CAD-CoASMCs, observing no significant increase compared to control cells (Fig. S1C). In a translational perspective, determining whether experimental models replicate these molecular changes is essential. To this end, we examined pACLY and ACLY expression levels in dissected PAs from rats exposed or not to Sugen/Hypoxia (Su/Hx), the best available model to study pulmonary hypertension associated with vascular remodeling (*14*). Western blot analysis revealed an augmentation in both active and total forms in the Su/Hx-exposed group (Fig. 1D). Furthermore, we demonstrated that ACLY expression is increased in experimental model of stenosis (e.g. denudation of the rat carotid artery with a flexible wire) that ACLY expression correlate with vascular remodeling in dogs subjected to coronary artery bypass grafting (CABG) (Fig. 1, E and F), providing additional evidence supporting the involvement of ACLY in adverse arterial remodeling.

**Figure 1.**
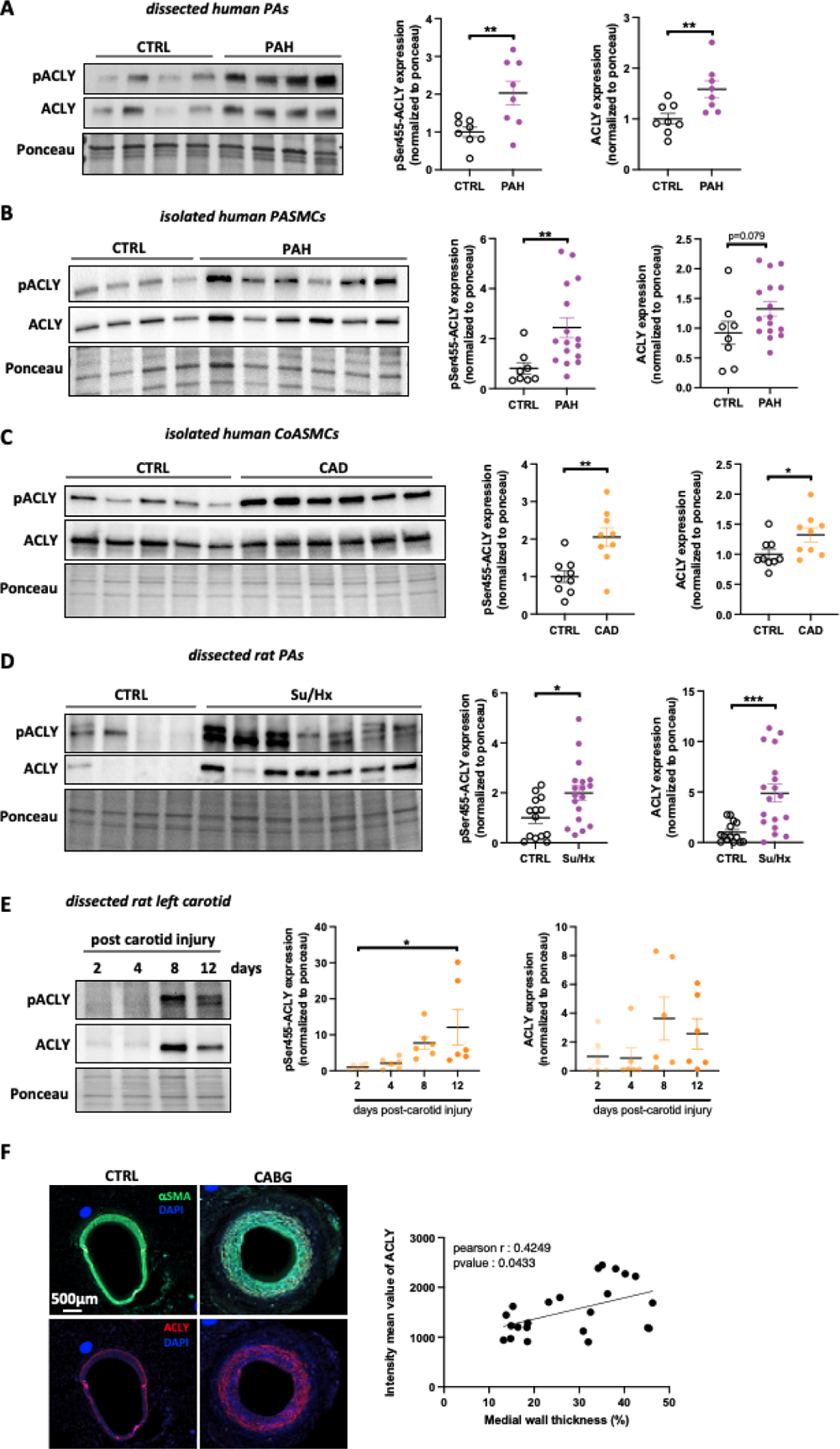
Elevated ACLY expression in remodeled arteries from PAH/CAD patients and animal models. **(A, B)** Representative Western blots and corresponding quantifications of phospho-ACLY (S455) and ACLY expression in dissected pulmonary arteries (PAs, A) and isolated PASMCs (B) from healthy donors and patients with PAH (n=8-16 **p<0.01, unpaired Student’s *t* test; data represent mean ± SEM). **(C)** Representative Western blots and corresponding quantifications of phospho-ACLY (S455) and ACLY in CoASMCs isolated from healthy donors and patients with CAD (n=9 *p<0.05, **p<0.01, unpaired Student’s *t* test; data represent mean ± SEM). **(D)** Representative Western blots and corresponding quantifications of phospho-ACLY (S455) and ACLY expression in dissected PAs from control and Sugen/Hypoxia (Su/Hx)-challenged rats (n=14 to 18; *p<0.05, ***p<0.01, unpaired Student’s *t* test; data represent mean ± SEM). **(E)** Representative Western blots and corresponding quantifications of phospho-ACLY (S455) and ACLY expression in carotid arteries from rats at 2, 4, 8, and 12 days after injury (n=6; *p<0.05, one-way ANOVA followed by Tukey’s post hoc analysis; data represent mean ± SEM). **(F)** Representative immunofluorescence images of αSMA and ACLY in saphenous veins from dogs after coronary arterial bypass grafting (CABG). Scale bar, 500μm. Correlation between ACLY fluorescence intensity and medial wall thickness (Pearson’s, r=0.4249; P<0.05).

### ACLY promotes PAH-PASMC and CAD-CoASMC survival, proliferation, and migration

To elucidate the functional significance of ACLY in fostering the pro-survival, pro-proliferative, and pro-migratory characteristics of PAH-PASMCs and CAD-CoASMCs, our initial investigation focused on the biological repercussions of silencing ACLY in diseased cells. Knockdown of ACLY using silencing RNAs (siRNA) exerted a noteworthy inhibitory effect on the proliferative capacity of both PAH-PASMCs and CAD-CoASMCs. This inhibition was evident through a significant decrease in the percentage of cells displaying nuclear Ki67 labeling and a concurrent reduction in the expression of cell proliferative markers such as Polo Like Kinase 1 (PLK1), Minichromosome Maintenance Complex Component 2 (MCM2) and proliferating cell nuclear antigen (PCNA) (Fig. 2, A to C). Furthermore, our findings revealed that ACLY knockdown significantly diminished the survival of PAH-PASMCs and CAD-CoASMCs. This was underscored by an elevated detection of Annexin V-positive cells and a concurrent reduction in the expression of Survivin (Fig. 2, A to C); which is documented to be highly implicated in the apoptosis resistance of SMCs in vascular diseases (*15*). Likewise, pharmacological inhibition of ACLY using BMS-303141 elicited comparable anti-survival and anti-proliferative effects in diseased cells (Fig. 2, A, D and E). To assess the impact of ACLY knockdown on the motility of diseased cells, a wound healing assay was conducted under serum-free conditions. In comparison to siSCRM-transfected cells, ACLY knockdown resulted in a notable reduction in the migration of both PAH-PASMCs and CAD-CoASMCs, although less pronounced in the latter (Fig. S2). Collectively, our results underscore the indispensable role of ACLY in driving the aberrant behavior of diseased SMCs, highlighting its significance in the pathological processes associated with PAH and CAD.

**Figure 2.**
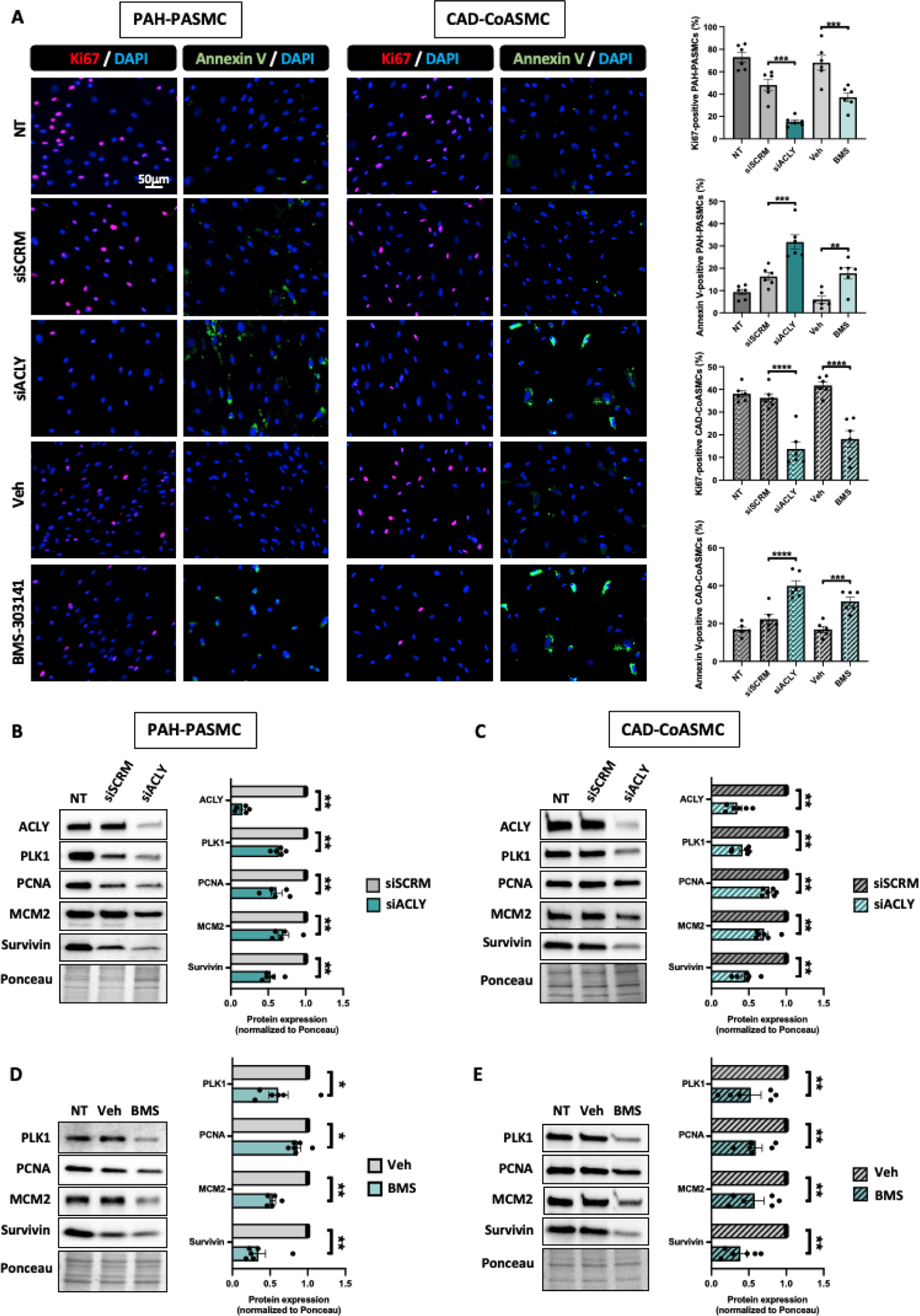
ACLY promotes PAH-PASMC and CAD-CoASMC proliferation and survival. **(A)** Representative fluorescent images, and corresponding quantifications of Ki67-labeled (red) and Annexin V-labeled (green) PAH-PASMCs and CAD-CoASMCs subjected or not to ACLY knockdown (siACLY) for 72h or exposed or not to BMS-303141 (30μM) for 48h (n=6; **p<0.01,***p<0.001, ****p<0.001 one-way ANOVA followed by Tukey’s post hoc analysis ; data represent mean ± SEM). Scale bar, 50μm. **(B and C)** Representative Western blots and corresponding quantifications of ACLY, PLK1, PCNA, MCM2, and Survivin expression in PAH-PASMCs (C) and CAD-CoASMCs (D) treated with siACLY or its negative control (siSCRM) for 72h (n=5 or 6; *p<0.05, **p<0.01,***p<0.001, Mann-Whitney’s test ; data represent mean ± SEM). **(D and E)** Representative Western blots and corresponding quantifications of PLK1, PCNA, MCM2, and Survivin expression in PAH-PASMCs (D) and CAD-CoASMCs (E) exposed or not to BMS-303141 (30μM) for 48h (n= 6; *p<0.05, **p<0.01, Mann-Whitney’s test; data represent mean ± SEM).

### Impact of molecular inhibition of ACLY on PAH-PASMC and CAD-CoASMC transcriptome

To comprehensively define the genes that are responsive to ACLY in diseased cells, we conducted RNA sequencing in 3 PAH-PASMC and 4 CAD-CoASMC cell lines subjected or not to ACLY knockdown by siRNA. Depletion of ACLY was confirmed by Western blot (Fig. S3A). In PAH-PASMCs, 1163 transcripts were found differentially expressed (absolute fold change of ≥1.5 and adjusted p-values ≤0.05), of which 634 were up-regulated and 529 were downregulated. Using the same threshold for expression changes and adjusted p-values, analysis revealed that 268 genes were up-regulated, and 313 genes were downregulated after depletion of ACLY in CAD-CoASMCs. The heatmap and volcano plots shown in Figure 3 highlight the genes significantly up- and down-regulated upon ACLY knockdown in diseased cells. As expected, ACLY was found strongly downregulated after siACLY transfection in both datasets, thus underscoring the significance of our RNA sequencing results (Fig. 3, A to D). To elucidate the functional implications of ACLY, we utilized ShinyGO v0.73.3 software to identify KEGG pathways influenced by ACLY knockdown in diseased cells. Notably, in both PAH-PASMCs and CAD-CoASMCs, gene signatures related to FA and cholesterol biosynthesis were markedly enriched among downregulated genes (Fig. 3, E and F). These findings align with established literature highlighting the crucial role of ACLY in *de novo* lipid synthesis and cell proliferation (*16*). Gene sets enriched among up-regulated genes were less conclusive. When comparing the modified sets of genes between both cell types, we found an overlap of 212 genes (88 up-regulated and 124 down-regulated, Fig. 3G). Gene ontology (GO) biological process analysis indicated that the overlapping downregulated genes were mainly enriched in cholesterol biosynthetic processes and cell division (Fig. 3H). Beyond ACLY itself, scrutiny of the genes constituting these signatures revealed downregulation of key factors involved in cholesterol biosynthesis, including 3-Hydroxy-3-Methylglutaryl-CoA Synthase 1 (HMGCS1) (*17*), Sterol regulatory element-binding protein-2 (SREBF2) (*18*), 24-Dehydrocholesterol Reductase (DHCR24) (36), and Mevalonate kinase (MVK) (*20*). Importantly, several genes known to promote SMCs survival and proliferation in the context of vascular diseases, such as Checkpoint kinase 1 (CHEK1) (19), Forkhead Box M1 (FOXM1) (*22*), Thymidine kinase 1 (TK1) (*23*) and S100 Calcium Binding Protein A4 (S100A4) (*24*), exhibited significant reduction in both cell types upon ACLY knockdown. The alterations in selected genes implicated in sterol biosynthesis and cell proliferation were further confirmed through quantitative RT-PCR experiments (Fig. S3B). In parallel, the impact of pharmacological ACLY inhibition using BMS-303141 was assessed, revealing a consistent decrease in the expression of the aforementioned genes, aligning with the RNA-Seq dataset. All tested genes exhibited similar expression changes upon exposure to BMS-303141 except for SREBF2 (Fig. S3B). These findings collectively underscore the intricate role of ACLY in shaping the genetic landscape and functional attributes of disease-associated SMCs.

**Figure 3.**
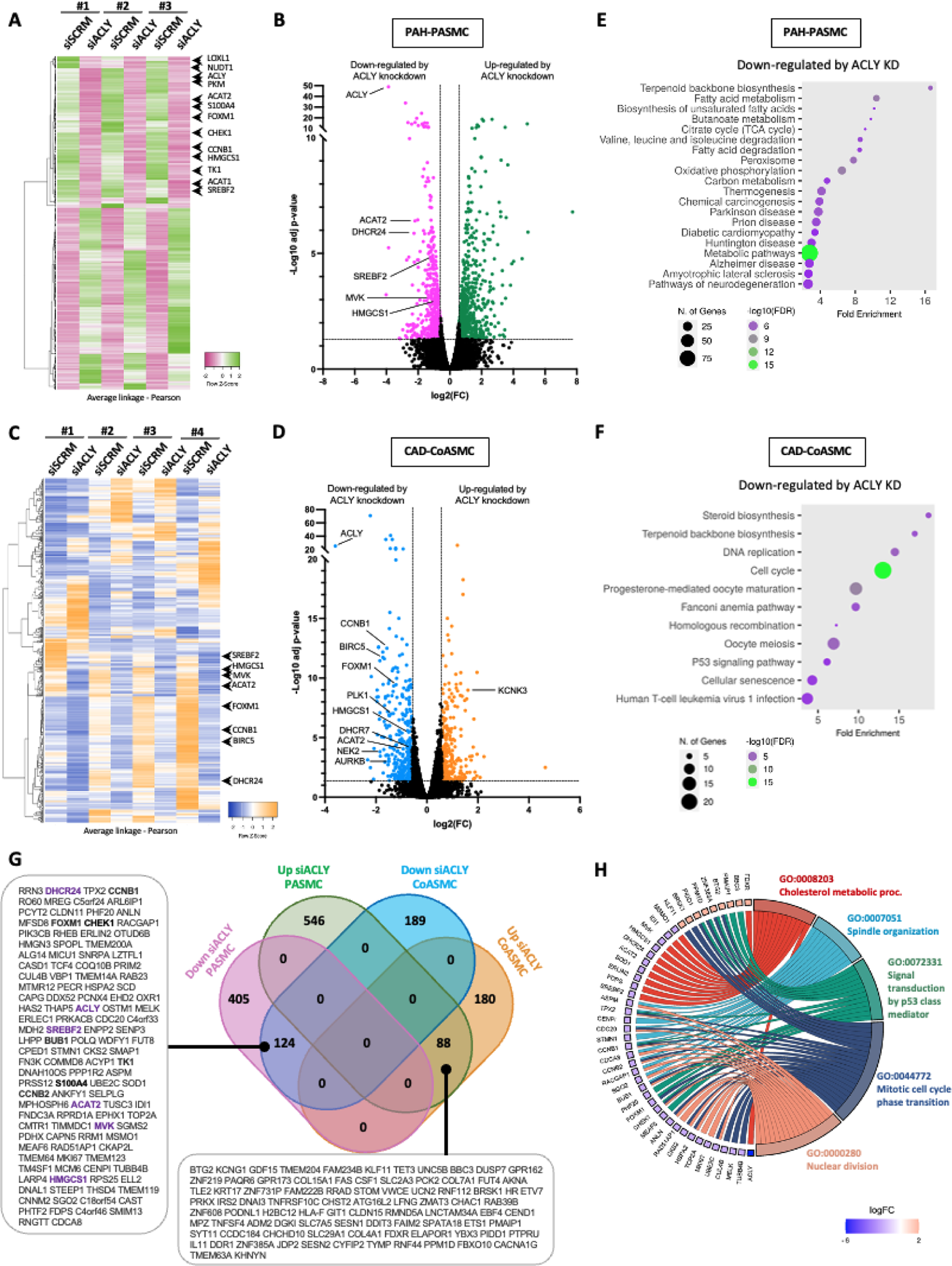
Transcriptomic response of diseased cells to ACLY knockdown. **(A)** Transcriptomic analysis of siRNA-mediated silencing of ACLY in PAH-PASMCs. Heatmap depicting the differentially expressed transcripts (|FC| ≥1.5 and adjusted p-values ≤0.05) as determined by RNA-seq analysis of siACLY- and siSCRM-treated PAH-PASMCs (n=3) for 72 hours. Expression values are depicted in line with the color scale; intensity increases from pink to green. Each column corresponds to a sample and each row indicates a transcript. **(B)** Volcano plot of log2 fold changes of all transcripts. The pink and green dots represent the significantly downregulated and upregulated transcripts in siACLY-treated vs siSCRM-treated PAH-PASMCs, respectively. **(C)** Transcriptomic analysis of siRNA-mediated silencing of ACLY in CAD-CoASMCs. Heatmap depicting the differentially expressed transcripts (|FC| ≥1.5 and adjusted p-values ≤0.05) as determined by RNA-seq analysis of siACLY- and siSCRM-treated CAD-CoASMCs (n=4) for 72 hours. Expression values are depicted in line with the color scale; intensity increases from blue to orange. Each column corresponds to a sample and each row indicates a transcript. **(D)** Volcano plot of log2 fold changes of all transcripts. The blue and orange dots represent the significantly downregulated and upregulated transcripts in siACLY-treated vs siSCRM-treated CAD-CoASMCs, respectively. **(E and F)** Enrichment analysis performed by ShinyGo software v0.77 showing the top biological processes (GO terms) in down-regulated genes by siACLY in PAH-PASMCs (E) and CAD-CoASMCs (F). Biological processes are ranked by fold enrichment values. The most significant processes are highlighted in green, and the less significant processes are highlighted in purple according to -Log10(FDR) values. Larger dots in the graph indicate a greater number of genes involved. **(G)** Venn diagrams showing the numbers of common and unique differentially expressed genes found to be significantly up- and down-regulated upon ACLY knockdown in both PAH-PASMCs and CAD-CoASMCs. **(H)** Chord plot showing the top 5 biological process (GO terms) enriched in overlapping down-regulated genes by siACLY. For left semicircle, the gradual color represents log fold change (FC) of each gene.

### ACLY inhibition impedes bioenergetics and lipogenesis

By actively consuming citrate and thus maintaining it at a low level, increased expression of ACLY was documented to sustain a glycolytic state in cancer cells (*8*). Given that a metabolic shift towards glycolytic metabolism is a characteristic of PAH-PASMCs, known to stimulate cell cycle progression, our initial investigation focused on the expression levels of various glycolytic enzymes in PAH-PASMCs subjected or not to ACLY inhibition. Treatment with BMS-303141 resulted in a decreased expression of Lactate dehydrogenase A (LDHA, which facilitates the transformation of pyruvate into lactate), a diminished ratio of phosphorylated Pyruvate dehydrogenase (PDH) to total PDH, and a reduced ratio of Pyruvate kinase isozymes M2 (PKM2) to M1 (PKM1) within the PAH-PASMCs (Fig. S4A). Additionally, a significant reduction of 6-Phosphofructo-2-Kinase/Fructose-2,6-Biphosphatase 3 (PFKBP3) was noted. Mirroring findings with PAH-PASMCs, ACLY inhibition led to lowered PFKFB3, LDHA and pPDH/PDH expression in CAD-CoASMCs (Fig. S4B). Conversely, in CAD-CoASMCs, treatment with BMS-303141 did not significantly affect the PKM2/PKM1 ratio.

To delve deeper into the role of ACLY in metabolism, we assessed the impact of ACLY inhibition on the metabolic parameters of PAH-PASMCs and CAD-CoASMCs. Specifically, we aimed to quantify changes in the oxygen consumption rate (OCR) and extracellular acidification rate (ECAR) with a Seahorse Extracellular Flux Analyzer. OCR serves as a measure of mitochondrial respiration, while ECAR reflects glycolysis. As expected, siACLY-treated PAH-PASMCs and CAD-CoASMCs exhibited increases in basal OCR, spare maximal OCR, respiratory capacity, and ATP production compared with siSCRM-transfected cells (Fig. S5, A to E). Concurrently, glycostress tests showed a decrease in glycolysis, glycolytic capacity, and glycolytic reserve in PAH-PASMCs and CAD-COASMCs treated with siACLY (Fig. S5, F to I). Interestingly, these findings illustrate that ACLY inhibition in both diseased cells markedly elevates the OCR/ECAR ratio (Fig. S5J), suggesting that metabolic energy is primarily derived from mitochondrial respiration at the expense of glycolysis.

Enhanced lipid synthesis is a well-documented metabolic trait of cancer (*25, 26*). As previously illustrated (Fig. 3G and S3B), depletion of ACLY was associated with a decreased expression of pivotal enzymes responsible for *de novo* cholesterol biosynthesis (DHCR24, SREBF2, and HMGCS1). To further elucidate the effects of ACLY knockdown on lipid metabolism, we quantified the expression of proteins involved in the *de novo* FA (Fig. 4A). Inhibition of ACLY was associated with an increase in the ratio of phosphorylated to total acetyl-CoA carboxylase (p-ACC/ACC) in PAH-PASMCs, signifying an increase in the inactive form of ACC, while a non-significant trend was observed in CAD-CoASMCs (Fig. 4B). Moreover, the expression levels of FA synthase (FASN) and Stearoyl-CoA desaturase (SCD), two additional enzymes crucial for FA synthesis, were diminished when ACLY expression was suppressed in both PAH-PASMCs and CAD-CoASMCs (Fig. 4B). At the cellular level, diverse staining techniques were employed to assess the impact of ACLY inhibition on cholesterol and FA synthesis in both PAH-PASMCs and CAD-CoASMCs. Filipin III, a polyene antibiotic that specifically binds free cholesterol, was utilized to gauge intracellular cholesterol levels. As expected, the suppression of ACLY resulted in a notable reduction in cholesterol accumulation in both diseased cells (Fig. 4C). Furthermore, Oil Red O staining, a fat-soluble dye to stain neutral lipids and cholesteryl esters (*27, 28*), revealed a diminished presence of intracellular lipid droplets, indicating lower lipid reserves (Fig. 4C). Cancer cells often sustain *de novo* synthesis of FAs as a protective mechanism to prevent excessive uptake of polyunsaturated FA (PUFAs), which can render the cells more susceptible to apoptosis through lipid peroxidation (*29*). In line with this, Xiang et al. (*30*) demonstrated that inhibition of ACLY promotes the accumulation of PUFAs (n-3 and n-6) in cancer cells through external absorption, inducing mitochondrial damages. Based on these findings, we postulated that ACLY inhibition would trigger intercellular PUFA accumulation in PAH-PASMCs and CAD-CoASMCs. Interestingly, mass spectrometry unveiled a marked rise in omega-3 (n-3) and n-6 PUFAs levels, including linoleic acide (LA) (18:2), eicosapentaenoic acid (EPA) (20:5), and docosahexaenoic acid (DHA) (22:6), in PAH-PASMCs when ACLY was deficient (Fig. S6). This suggests heightened external PUFA uptake. While a comparable trend was noted in CAD-CoASMC, statistical significance was not attained.

**Figure 4.**
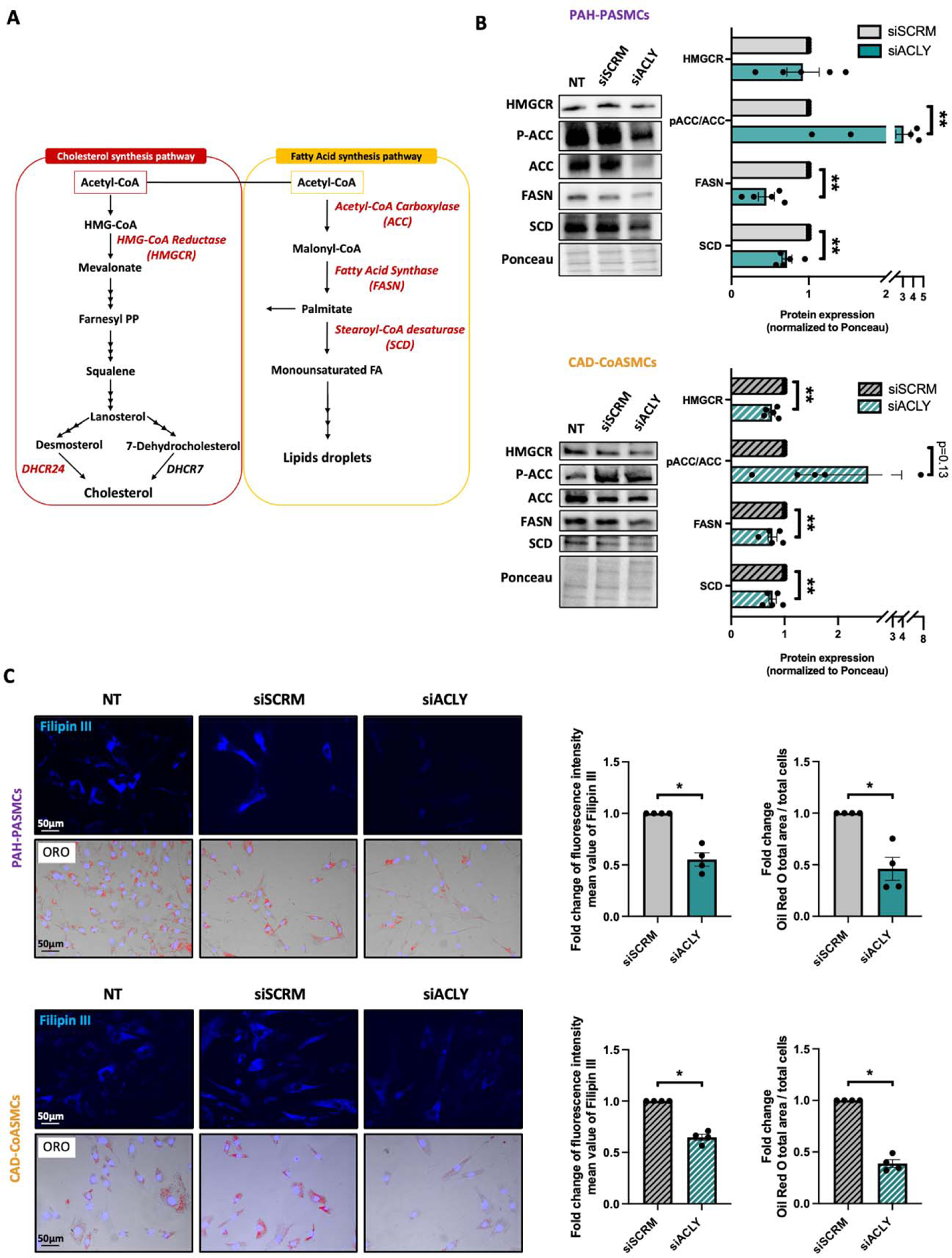
ACLY promotes PAH-PASMC and CAD-CoASMC de Novo lipid and synthesis. **(A)** Representative fluorescent images, and corresponding quantifications of filippin-3 labeled (blue) and oil-red o (red) PAH-PASMCs and CAD-CoASMCs subjected or not to ACLY knockdown (siACLY) for 72h. (n=4; *p<0.05, Mann-Whitney’s test; data represent mean ± SEM). Scale bar, 50μm. **(B)** Simplified schematic of cholesterol and fatty acid synthesis pathways. **(C)** Representative Western Blots and corresponding quantification of HMGCR, phospho-ACC (S79), ACC, FASN and SCD expression in PAH-PASMCs and CAD-CoASMCs subjected or not to ACLY knockdown using siRNA for 72h (n=5; **p<0.01, Mann-Whitney’s test; data represent mean ± SEM).

Taken together, our results suggest that silencing ACLY in PAH-PASMCs and CAD-CoASMCs disrupts lipid metabolism pathways, leading to decreased cholesterol and FA synthesis, and promoting the accumulation of PUFAs which could render cells more susceptible to apoptosis(*30*).

### ACLY increases nuclear acetyl-CoA levels and promotes epigenetic reprogramming of proliferative genes

Considering that ACLY is described as a nucleo-cytoplasmic enzyme responsible for producing acetyl-CoA crucial for histone acetylation (*31*), we then determined if differences in ACLY protein expression between control and PAH-PASMCs are accompanied by changes in its subcellular localization. To investigate this, we employed immunofluorescence analysis and found that ACLY was more intensely labeled in the nuclei of PAH-PASMCs compared to their normal counterparts (Fig. 5A). A similar result was observed in CAD-CoASMCs (Fig S7A). To further support our findings, we employed a subcellular fractionation approach followed by Western blotting to compare ACLY expression in cytosolic and nuclear fractions from control and diseased cells. Histone H3 and α/β Tubulin served as loading controls for nuclear and cytosolic fractions, respectively. As depicted in Figure 5B and Figure S7B, ACLY expression was more pronounced in the nucleus of PAH-PASMCs and CAD-CoASMCs than in control cells, suggesting that ACLY contributes to the nuclear acetylation process and transcriptional changes in diseased cells. To further investigate this point, we measured the amount of acetyl-CoA in nuclear fractions of PAH-PASMCs or CAD-CoASMCs, with or without ACLY depletion, revealing decreased levels of nuclear acetyl-CoA in siACLY-transfected cells (Fig. 5C). This supports the role of ACLY in supplying nuclear acetyl-CoA. Accordingly, levels of acetylated histones (H3K27, H3K9, and H4) were lower in PAH-PASMCs and CAD-CoASMCs upon ACLY silencing (Fig 5, D and E).

**Figure 5.**
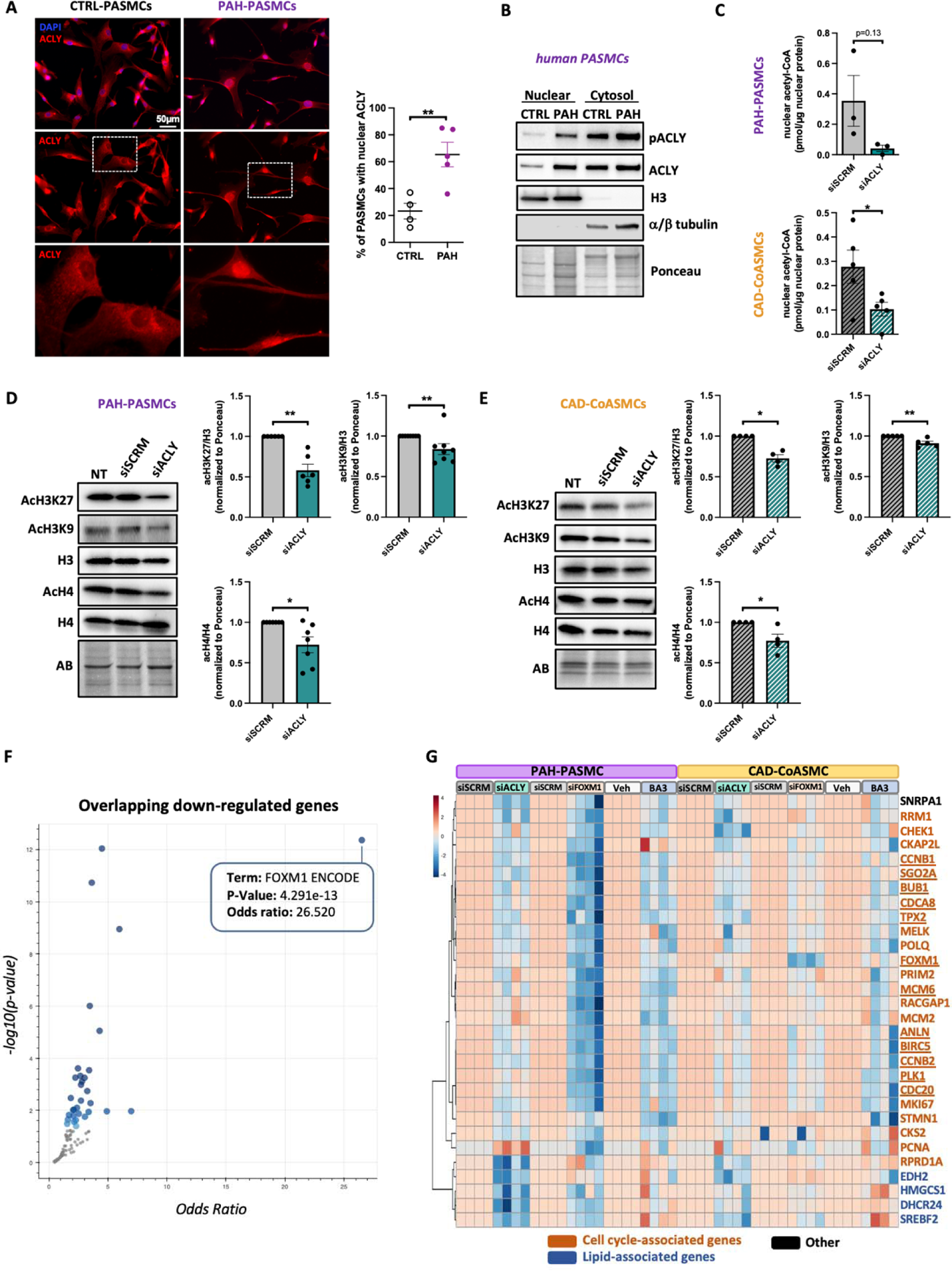
ACLY regulates histone acetylation and FOXM1 activation. **(A**) Representative fluorescent images of control (n=4) and PAH-PASMCs (n=5) labeled with ACLY. Percentage of cells exhibiting nuclear localization of ACLY is shown (n=4 or 5; **p<0.01, ***p<0.001 unpaired Student’s *t* test; data represent mean ± SEM). Scale bar, 50μm. **(B)** Representative Western blots (of 3 independent experiments) for phospho-ACLY (S455), ACLY, α/β tubulin, and histone H3 (H3) in nuclear and cytosolic extracts from control and PAH-PASMCs. **(C)** Acetyl-CoA levels measured in nuclear fractions of PAH-PASMCs and CAD-CoASMCs exposed or not to siACLY for 72h (n=3 to 5; *p<0.05, unpaired Student’s *t* test; data represent mean ± SEM). **(D)** Representative Western blots and corresponding quantifications of AcH3K27, AcH3K9, H3, AcH4 and H4 expression in PAH-PASMCs treated with siACLY or its negative control (siSCRM) for 72h (n=6 or 7; *p<0.05, **p<0.01 Mann-Whitney’s test; data represent mean ± SEM). **(E)** Representative Western blots and corresponding quantifications of AcH3K27, AcH3K9, H3, AcH4 and H4 expression in CAD-CoASMCs treated with siACLY or its negative control (siSCRM) for 72h (n=4; *p<0.05, **p<0.01 Mann-Whitney’s test; data represent mean ± SEM). **(F)** Volcano plot of terms from the encode and ChEA consensus TFs from ChIP-X gene set. Each point represents a single term, plotted by corresponding odds ratio (x-position) and -log10 (p-value) (y-position) from the enrichment results of the input query gen set. The larger and darker-colored the point, the more significantly enriched the input gene set is for the term. **(G)** Fold change heatmap showing the qPCR analysis results of selected genes under siACLY; siFOXM1 (72h) or Butyrolactone 3 (BA3, 70μM, 48h) treatments in PAH-PASMCs and CAD-CoASMCs. The 12 underlined genes are the downregulated genes in both diseases by ACLY, FOXM1 and GCN5 inhibition. Fold change calculate based on delta Ct value compared to the control samples, red implies increased expression while blue implies decreased expression. (n=4 or 6; *p<0.05, **p<0.01 Mann-Whitney’s test).

Interrogation of the chromatin immunoprecipitation enrichment analysis (ChEA) 2022 gene set library using the EnrichR web server revealed FOXM1 as the top enriched transcription factor governing the expression of the 124 shared downregulated genes across both cell types (Fig. 5F). Of note, FOXM1 was part of these genes and has been confirmed to be downregulated upon ACLY inhibition (Fig. 3G; S3 and S7C). Based on these data, we hypothesized that the downregulation of genes resulting from ACLY knockdown could be attributed to the inhibition of the transcription factor FOXM1. To validate this hypothesis, we conducted qRT-PCR on 30 genes, all known for their association with proliferation and lipid metabolism, out of the 124 commonly downregulated genes in both CAD and PAH. Our results demonstrated that among the 30 genes, 26 in PAH-PASMCs and 24 in CAD-CoASMCs were significantly downregulated by siACLY, as confirmed by qRT-PCR (Table S1). Similarly, employing siFOXM1, we observed downregulation of 25 out of 30 genes in PAH-PASMCs and 22 out of 30 genes in CAD-CoASMCs, including FOXM1 itself (Table S1). These results provide strong support for our hypothesis, suggesting that the reduction in gene expression observed after ACLY knockdown is likely a result of FOXM1 inhibition.

In terms of mechanisms, acetylation of FOXO1 at lysine 294 (Lys294) was shown to reduce its affinity for DNA binding(*32*), favoring the action of its natural antagonist FOXM1 which competes for the same gene targets (*33*). Consistently, we found that ACLY inhibition, by reducing acetyl-CoA generation, leads to decreased acetylation in both PAH-PASMCs and CAD-CoASMCs (Fig. S7C), ultimately resulting in the repression of FOXM1.

In addition, and likely in combination with FOXM1, the decrease in gene expression caused by ACLY knockdown is expected to result from a decrease in activity of histone acetyltransferases (HATs). HATs function as enzymes that transfer an acetyl group from acetyl-CoA to specific histone proteins, leading to their acetylation. This acetylation process is vital for enhancing gene expression. Considering that ACLY is known to overlap with the histone acetylation targets of GCN5 in various cell types(*9, 34*) and GCN5 is reported to promote cell proliferation and adipogenesis (*35, 36*), we investigated whether ACLY signature on genes expression could be mediated by GCN5-histone acetylation. We initially assessed the expression levels of GCN5 in control versus PAH-PASMCs and control versus CAD-CoASMCs. GCN5 exhibited a significant increase in both diseased cells (Fig. S8A). We next explored the impact of pharmacological GCN5 inhibition using Butyrolactone (BA3) on the pro-proliferative and apoptosis-resistance phenotype of PAH-PASMCs and CAD-CoASMCs. Initially, we validated the influence of BA3 on GCN5 activity, noting a reduction in the histone acetylation profile (Fig S8B). Pharmacological inhibition of GCN5 replicated the anti-survival and anti-proliferative effects observed with ACLY inhibition (Fig. S8, C and D). Moreover, the inhibition of GCN5 in PAH-PASMCs and CAD-CoASMCs leads to a reduction in FOXM1 expression, which is similar to the effects observed with ACLY knockdown (Fig. S9A). While BA3 is highly specific to GCN5, it is noteworthy that it can also inhibit PCAF activity, another HAT highly similar to GCN5. We assessed FOXM1 expression levels in PAH-PASMCs and CAD-CoASMCs upon siRNA-mediated PCAF knockdown. No difference was observed, demonstrating that BA3’s effects are attributable to GCN5 inhibition (Fig. S9B). We then measured the effects of GCN5 inhibition on the expression of the 30 genes selected from the common set of 124 genes, similarly to what was done with ACLY and FOXM1 knockdown. We observed that among these 30 shared genes, 16 were significantly downregulated by BA3 in both CAD and PAH conditions (Table S1). A total of 12 genes showed similar downregulation upon inhibition of ACLY, FOXM1, and GCN5 in both cell types. Interestingly, among these 12 genes, 11 displayed increased expression levels in the disease group compared to the control group in both diseased cells (Table S1). These 12 genes are primarily associated with cell cycle and proliferation processes including CCNB1, CCNB2, FOXM1, PLK1 and survivin.

Overall, these results collectively provide robust evidence supporting the hypothesis that the downregulation of the investigated genes upon ACLY inhibition is regulated at least in part by GCN5 and FOXM1.

### Pharmacological inhibition of ACLY using BMS-303141 improves established PAH in Sugen/Hypoxia-challenged rats

To stress the importance of ACLY in PAH progression, BMS-303141 was administered for two weeks to Su/Hx rat after PAH was established (Fig. 6A). As indicated by right heart catheterization (RHC), treatment with BMS-303141 resulted in significant diminution of right ventricular systolic pressure (RVSP) and mean PA pressure (mPAP) compared with the vehicle-treated Su/Hx rats (Fig. 6B and Fig S10A). Stroke volume (SV) and cardiac output (CO) were partially recovered by BMS-303141 administration (Fig. 6B and Fig. S10A). As predicted by hemodynamic data, pulmonary vascular remodeling was improved by BMS-303141, as indicated by reduced total pulmonary resistance (TPR) and morphometric analysis showing decreased wall thickness of small PAs (Fig. 6D and Fig. S10A). In agreement with this, the media of the remodeled Su/Hx PAs showed evidence of enhanced cell proliferation, with increased PCNA-positive PASMCs, whereas in BMS-303141-treated animals, the percentage of PASMCs-positive for PCNA was significantly lower (Fig. 6E and Fig. S10B). Conversely, there was a significant increase in the percentage of apoptotic PASMCs, as revealed using immunofluorescence for cleaved caspase-3, in BMS-303141-treated rats (Fig. 6F and Fig. S10B). This dual effect on cell proliferation and apoptosis underscores the therapeutic potential of targeting ACLY in mitigating PAH progression. Moreover, immunofluorescent staining indicated a diminution in the acetylation of lysine 27 and lysine 9 on histone H3 (H3K27 and H3K9, respectively) in PASMCs from BMS-303141-treated Su/Hx animals in comparison to those administered with vehicle (Fig. 6G and Fig. S10B). This reduction in histone acetylation may reflect a global reduction in transcriptional activation since acetylation at these residues is typically associated with chromatin de-condensation and increased gene expression.

**Figure 6.**
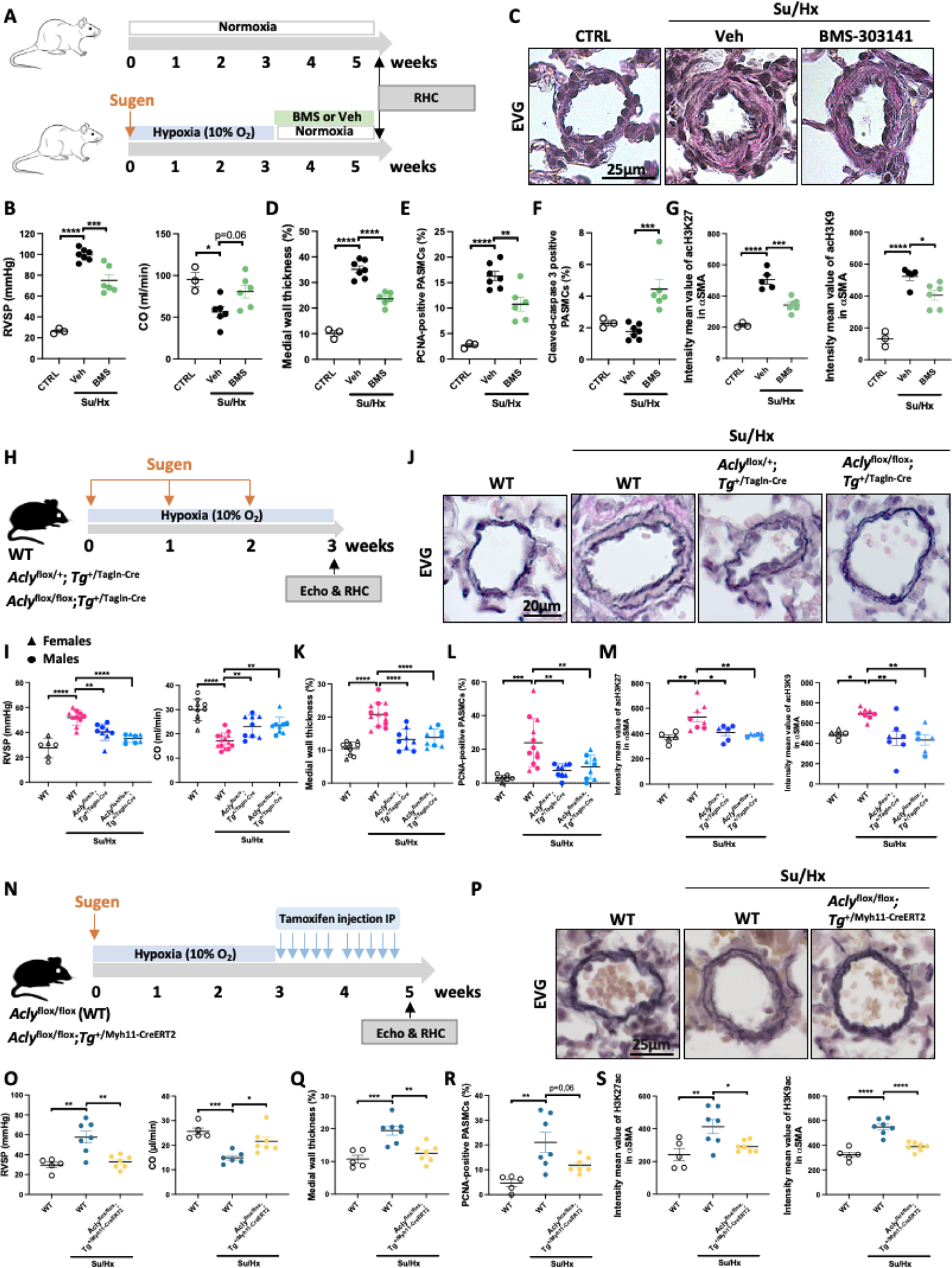
Pharmacological inhibition of ACLY and *Acly* loss-of-function targeted to smooth muscle cells mitigate PAH in Sugen/Hypoxia rats. **(A)** Experimental design whereby Sprague Dawley rats were injected with SU5416 (Su) and exposed to chronic hypoxia (Hx) for 3 weeks to induce PAH. Subsequently, PAH rats were treated with BMS-303141 versus vehicle for 2 weeks. **(B)** RVSP and CO in BMS-303141-treated Su/Hx PAH rats versus controls as assessed by right heart catheterization (n=3 to 7; *p<0.05,***p<0.001, ****p<0.001one-way ANOVA followed by Tukey’s post hoc analysis; data represent mean ± SEM). **(C)** Representative images of distal PAs stained with Elastica van Gieson (EVG). Scale bar, 25μm. **(D)** Quantification of medial wall thickness in control, Su/Hx+Veh and Su/Hx+BMS rats (n=3 to 7; ****p<0.001, one-way ANOVA followed by Tukey’s post hoc analysis; data represent mean ± SEM). **(E)** Immunofluorescent quantifications for PCNA (ANOVA followed by Tukey’s post hoc analysis) **(F)**, cleaved caspase-3 Kruskal-Wallis followed by Dunn’s post hoc analysis (n=3 to 7) **(G),** acH3K27 and acH3K9 (one-way ANOVA followed by Tukey’s post hoc analysis) in BMS-303141-treated Su/Hx PAH rats versus controls (n=3 to 7; *p<0.05 ** p<0.01***p<0.001, ****p<0.0001 data represent mean ± SEM). **(H)** Schematic of Su/Hx protocol in wildtype (WT), *Acly*^+/flox^;Tg^+/Tagln-Cre^, and *Acly*^flox/flox^;Tg^+/Tagln-Cre^ male and female mice. **(I)** RVSP measured by right heart catheterization and CO measured by echocardiography in Su/Hx-exposed WT, *Acly*^+/flox^;Tg^+/Tagln-Cre^, and *Acly*^flox/flox^;Tg^+/Tagln-Cre^ mice vs controls. (n=6 to 13; *p<0.05,**p<0.01, ****p<0.0001 (one-way ANOVA followed by Tukey’s post hoc analysis; data represent mean ± SEM). **(J)** Representative images of distal PAs stained with EVG. Scale bar, 20μm. **(K)** Quantification of medial wall thickness of distal PAs in control, Su/Hx-exposed WT, *Acly*^+/flox^;Tg^+/Tagln-Cre^, and *Acly*^flox/flox^;Tg^+/Tagln-Cre^ mice (n=9 to 13; ****p<0.0001, one-way ANOVA followed by Tukey’s post hoc analysis data represent mean ± SEM). **(L** and **M)** Immunofluorescent quantifications for PCNA **(L),** acH3K27 and acH3K9 **(M)** in distal PAs of control, Su/Hx-exposed WT, Acly^+/flox^_;_Tg^+/Tagln-Cre^, and Acly^flox/flox^;Tg^+/Tagln-Cre^ mice (n=7 to 12;, *p<0.05,**p<0.001,***p<0.001, one-way ANOVA followed by Tukey’s post hoc analysis; data represent mean ± SEM). **(N)** Experimental design whereby male WT and *Acly*^flox/flox^;Tg^+/Myh11-CreERT2^ mice were exposed to Su/Hx for 3 weeks to induce PAH followed by treatment with Tamoxifen for 2 weeks to induce *Acly* deletion targeted to smooth muscle cells. **(O)** RVSP and CO in Su/Hx-exposed WT and *Acly*^flox/flox^;Tg^+/Myh11-CreERT2^ mice vs controls. (n=5 to 7; *p<0.05, **p<0.01, ***p<0.001 one-way ANOVA followed by Tukey’s post hoc analysis; data represent mean ± SEM). **(P)** Representative images of distal PAs stained with EVG. Scale bar, 25[m. **(Q)** Quantification of medial wall thickness of distal PAs in control, Su/Hx-exposed WT and *Acly*^flox/flox^;Tg^+/Myh11-CreERT2^ mice (n=5 to 7; **p<0.01, ***p<0.001 one-way ANOVA followed by Tukey’s post hoc analysis data represent mean ± SEM). **(R** and **S)** Immunofluorescent quantifications for PCNA **(R),** acH3K27 and acH3K9 **(S)** in distal PAs of control, Su/Hx-exposed WT and *Acly*^flox/flox^;Tg^+/Myh11-CreERT2^ mice vs controls. (n=5 to 7; *p<0.05, **p<0.01, ***p<0.001,****p<0.0001 one-way ANOVA followed by Tukey’s post hoc analysis; data represent mean ± SEM).

### Smooth muscle-specific inactivation of *Acly* attenuates PAH development in Sugen/Hypoxia-challenged mice

An inherent concern with the use of pharmacological inhibitors is their specificity. Thus, to prove the pivotal role of smooth muscle ACLY in PAH development and progression and circumvent the embryonic lethality associated with its complete somatic disruption (55), adult SMC-specific *Acly* homozygous (*Acly*^flox/flox^;Tg^+/Tagln-Cre^) and heterozygous (*Acly*^+/flox^;Tg^+/Tagln-Cre^) mice were generated and injected once weekly with Sugen and exposed to chronic hypoxia for 3 weeks (Fig. 6H). Efficiency of *Acly* deletion was confirmed by Western blot since ACLY was nearly absent in pooled dissected PAs from the *Acly*^flox/flox^;Tg^+/Tagln-Cre^ mice (Fig. S11A). Echocardiographic assessment validated that PAH was present in Su/Hx-exposed control (*Acly*^flox/flox^) mice, as evidence by their decreased PA acceleration time (PAAT) when compared to nontreated control mice (Fig. S11B). When compared to Su/Hx control mice, *Acly*^flox/flox^;Tg^+/Tagln-Cre^ and *Acly*^+/flox^;Tg^+/Tagln-Cre^ mice subjected to Su/Hx exhibited increased PAAT, tricuspid annular plane systolic excursion (TAPSE), S wave, SV and CO, indicating decreased PAH severity (Fig. 6I and Fig, S11B). In line with data from echocardiography analysis, the elevation of RVSP and mPAP, measured by RHC, was attenuated in Su/Hx-subjected *Acly*^flox/flox^;Tg^+/Tagln-Cre^ and *Acly*^+/flox^;Tg^+/Tagln-Cre^ mice (Fig. 6I and Fig, S11C). Further examination revealed that Su/Hx-exposed *Acly*^flox/flox^;Tg^+/Tagln-Cre^ and *Acly*^+/flox^;Tg^+/Tagln-Cre^ mice displayed significant decreased medial wall thickness, fewer proliferating PASMCs and reduction of H3K27 and H3K9 acetylation as compared to Su/Hx WT mice ((Fig. 6, J to M) and Fig. S11D).

As a complementary approach, we evaluated the therapeutic benefit of *Acly* deletion on disease progression. To this end, mice expressing a tamoxifen-inducible Cre recombinase under the control of the SMC-specific promoter, *Myh11,* were used to generate *Acly*^flox/flox^;*Tg*^+/Myh11-CreERT2^ mice. After Su/Hx-induced PAH, mice were intraperitoneally injected with Tamoxifen and sacrificed after 2 weeks, according to the protocol illustrated in Figure 6N. Efficiency of *Acly* deletion was confirmed by qPCR using pooled aorta from several mice (Fig S12A). Echocardiographic assessment demonstrated that *Acly* inactivation resulted in an improvement of cardiac function, as illustrated by an increase in PAAT, TAPSE, S waves, and CO (Fig. 6O and Fig, S12B). RVSP and mPAP, as assessed by RHC, were significantly decreased (Fig. 6O and Fig. S12C). These hemodynamic improvements were associated with an attenuation of medial wall thickness (Fig. 6, P and Q). Moreover, the quantification of PCNA-positive PASMCs and the assessment of H3K27 and H3K9 acetylation showed a discernible decrease in these parameters (Fig. 6, R and S; and Fig S12D). Collectively, our results provide direct genetic evidence that ACLY expression in SMCs drives pulmonary vascular remodeling.

### Pharmacological and genetic inhibition of ACLY attenuates carotid stenosis *in vivo*

Based on the aforementioned results obtained from PAH models, we next investigated whether ACLY inhibition may also represent a therapeutic target in the context of stenosis. To address this question, carotid injury was induced in adult rats by mechanical denudation of the vascular endothelium followed by systemic administration of BMS-303141 or its vehicle for two weeks (Fig. 7). Two weeks after injury, a reproducible and severe intimal thickening was apparent after injury when compared with sham-operated animals (Fig. 7, A and B). Pharmacological inhibition of ACLY notably repressed neointimal thickening as manifested by a significant decrease in the intimal area and intima/media area ratio (I/M ratio) (Fig. 7C). This was accompanied by a significant reduction of the luminal obliteration (Fig. 7C). Accordingly, immunofluorescence showed that inhibition of ACLY mitigated the expression of PCNA, acH3K27 and acH3K9 in intimal cells of injured carotid arteries (Fig. 7, D and E; and Fig. S13).

**Figure 7.**
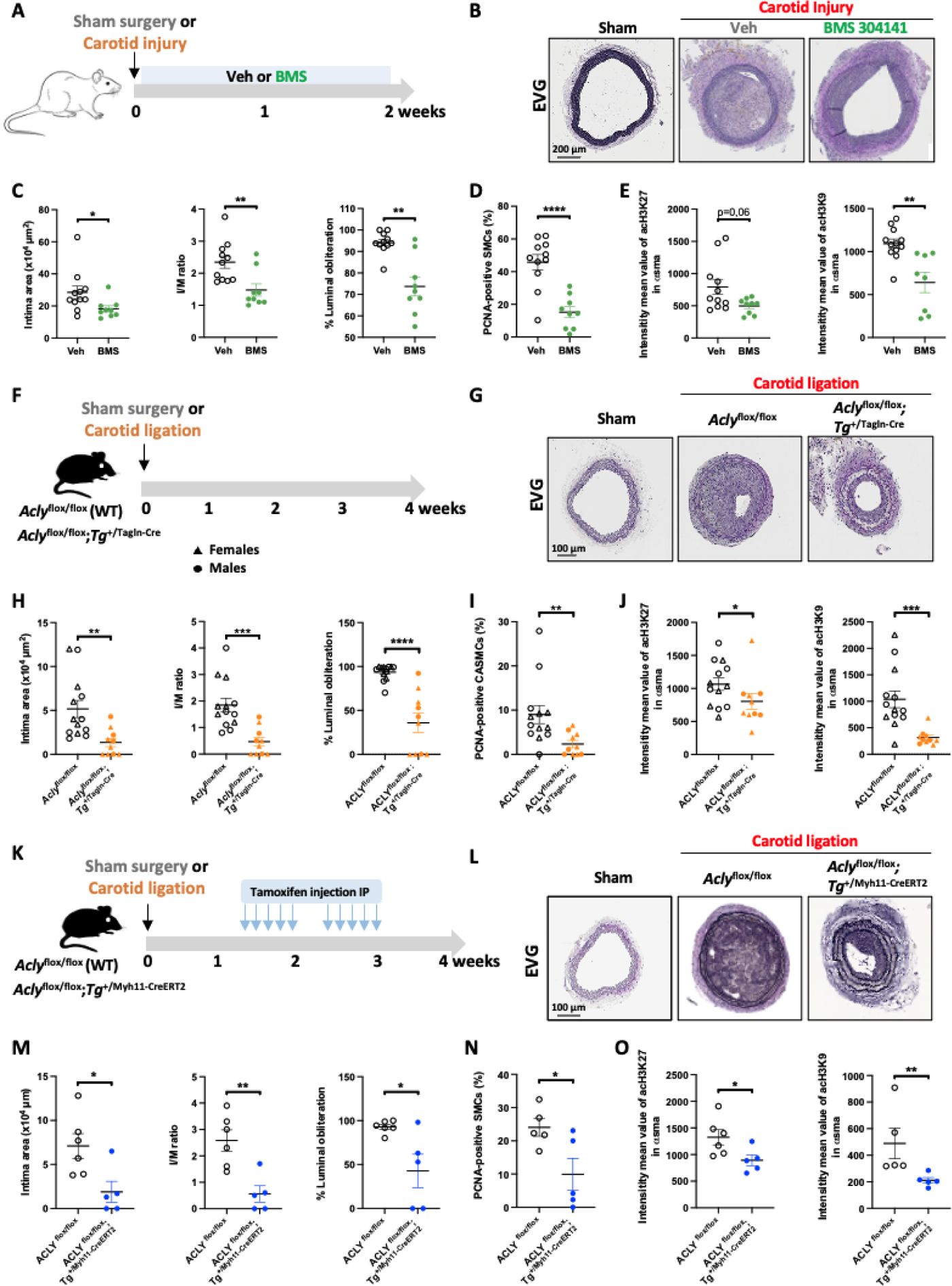
Pharmacological inhibition of ACLY and *Acly* loss-of-function targeted to smooth improve carotid remodeling induced by injury and ligation. **(A**) Schematic diagram of the design of the carotid artery injury experiment in Sprague Dawley rats. **(B)** Representative images of the carotid artery stained with Elastica van Gieson (EVG) in rats subjected or not to carotid injury and treated or not with BMS-303141 for 2 weeks. Scale bar, 200μm. **(C)** Quantifications of intimal area, intimal to media ratio (I/M), and luminal obliteration at 2 weeks after vascular injury (n=9 to 11; *p<0.05, **p<0.01, Mann-Whitney’s test; data represent mean ± SEM). **(D-E)** Immunofluorescent quantifications for PCNA **(D)** (unpaired Student’s *t* test, ****p<0.0001), acH3K27 (Mann-Whitney’s test p=0.06) and acH3K9 (unpaired Student’s *t* test **p<0.01) **(E)** in carotid arteries of rats subjected to vascular injury and treated or not with BMS-303141 for 2 weeks. (n=9 to 11; data represent mean ± SEM). **(F)** Schematic of carotid ligation protocol in wildtype (WT, *Acly*^flox/flox^) and *Acly*^flox/flox^;Tg^+/Tagln-Cre^ male and female mice. **(G)** Representative images of the carotid artery stained with EVG in WT and *Acly*^flox/flox^;Tg^+/Tagln-Cre^ mice subjected or not to carotid ligation at week 4. Scale bar, 100μm. **(H)** Quantifications of intimal area (Mann-Whitney’s test), I/M (unpaired Student’s *t* test), and luminal obliteration (Mann-Whitney’s test) in WT and *Acly*^flox/flox^;Tg^+/Tagln-Cre^ mice subjected to carotid ligation at week 4 (n=10 to 13; **p<0.01, ***p<0.001, ****p<0.0001; data represent mean ± SEM). **(I** and **J)** Immunofluorescent quantifications for PCNA **(I),** acH3K27 and acH3K9 **(J)** in carotid arteries of WT and *Acly*^flox/flox^;Tg^+/Tagln-Cre^ mice subjected to vascular injury (n=10 to 13; *p<0.05, **p<0.01, ***p<0.001, Mann-Whitney’s test data represent mean ± SEM). **(K)** Schematic of carotid ligation protocol in wildtype (WT, *Acly*^flox/flox^) and *Acly*^flox/flox^;Tg^+/Myh11-CreERT2^ male mice. Mice were injected with Tamoxifen to induce *Acly* deletion targeted to smooth muscle cells post vascular injury. **(L)** Representative images of the carotid artery stained with EVG in Tamoxifen-treated WT and *Acly*^flox/flox^;Tg^+/Myh11-CreERT2^ mice subjected or not to carotid ligation at week 4. Scale bar, 100[m. **(M)** Quantifications of intimal area, I/M, and luminal obliteration in Tamoxifen-treated WT and *Acly*^flox/flox^;Tg^+/Myh11-CreERT2^ mice subjected to carotid ligation at week 4. (n=5 or 6; *p<0.05, **p<0.01, unpaired Student’s *t* test; data represent mean ± SEM). **(N** and **O)** Immunofluorescent quantifications for PCNA **(N),** acH3K27 and acH3K9 **(O)** in Tamoxifen-treated WT and *Acly*^flox/flox^;Tg^+/Myh11-CreERT2^ mice subjected to carotid ligation at week 4. (n=5 or 6; *p<0.05, **p<0.01, unpaired Student’s *t* test; data represent mean ± SEM).

To further demonstrate the crucial requirement of ACLY in response to vascular injury, 8-10-week-old *Acly*^flox/flox^;Tg^+/Tagln-Cre^ mice and their WT littermates were subjected to carotid artery ligation (Fig. 7F). Immunostaining confirmed reduced expression of ACLY in *Acly*^flox/flox^;Tg^+/Tagln-Cre^ mice (Fig. S14A). The time course of carotid injury demonstrated the complete obliteration of the vessels 28 days post-injury (Fig. S14B). Thus, mice were euthanized on day 28 and left carotid arteries were harvested. Histomorphometric analysis showed that *Acly* loss-of-function targeted to SMCs significantly attenuated carotid artery injury-induced increases in neointimal area, I/M ratio, and luminal obliteration (Fig. 7H). Additionally, intimal PCNA-positive cells were significantly reduced in injured *Acly*^flox/flox^;Tg^+/Tagln-Cre^ mice (Fig. 7I and Fig. S14C). Similarly, acetylation levels of H3K27 and H3K9 were reduced in mice with *Acly* deletion (Fig. 7J and Fig. S14C).

As a complementary approach, we evaluated the therapeutic benefit of ACLY inhibition on disease progression. To this end, *Acly^f^*^lox/flox^ mice were bred with *Tg*^+/Myh11-CreERT2^ mice. Because the time course of carotid injury revealed the initiation of neointimal hyperplasia 7 days following carotid artery ligation (Fig. S14B), tamoxifen injection was initiated at day 10 and *Acly^f^*^lox/flox^ (WT) and *Acly*^flox/flox^;*Tg*^+/Myh11-CreERT2^ mice were sacrificed at day 28, according to the protocol depicted in Figure 7K. Efficiency of *Acly* deletion was confirmed by qPCR (Fig S12A). Analysis showed that deletion of *Acly* significantly attenuated carotid injury-induced intimal hyperplasia compared with that in WT mice, as demonstrated by a decreased intimal area, I/M ratio, and lumen obliteration (Fig. 7, L and M) along with a reduction in PCNA-positive cells and histone acetylation (Fig 7, N and O; and Fig. S15).

Taken together, our findings indicate that ACLY is an essential component of the signaling cascade resulting in neointima formation after carotid artery injury.

Inhibition of ACLY using BMS-303141 reduces vessel remodeling in *ex vivo* human precision cut lung slices, human coronary arteries and human saphenous vein rings Precision-cut lung slices (PCLS) are an innovative experimental model that offer a unique approach to studying lung physiology, pharmacology, and pathophysiology within an organotypic context. PCLS supports studies on live tissue for several days, enabling the examination of both acute and chronic effects of drugs, thereby increasing the translational potential of the research findings. In our model, PCLS were exposed to a combination of growth factors (endotheline-1, PDGF-BB and FGF2) to induce vessel wall remodeling (Fig. 8A)(*38*). As expected, PAs exposed to growth factors exhibited notable medial wall thickening compared to untreated PAs (Fig. 8B). In all biologically distinct samples, ACLY inhibition using BMS-303141 significantly decreased medial wall thickness, PASMCs survival (C3C) and proliferation (PCNA) (Fig. 8B).

**Figure 8.**
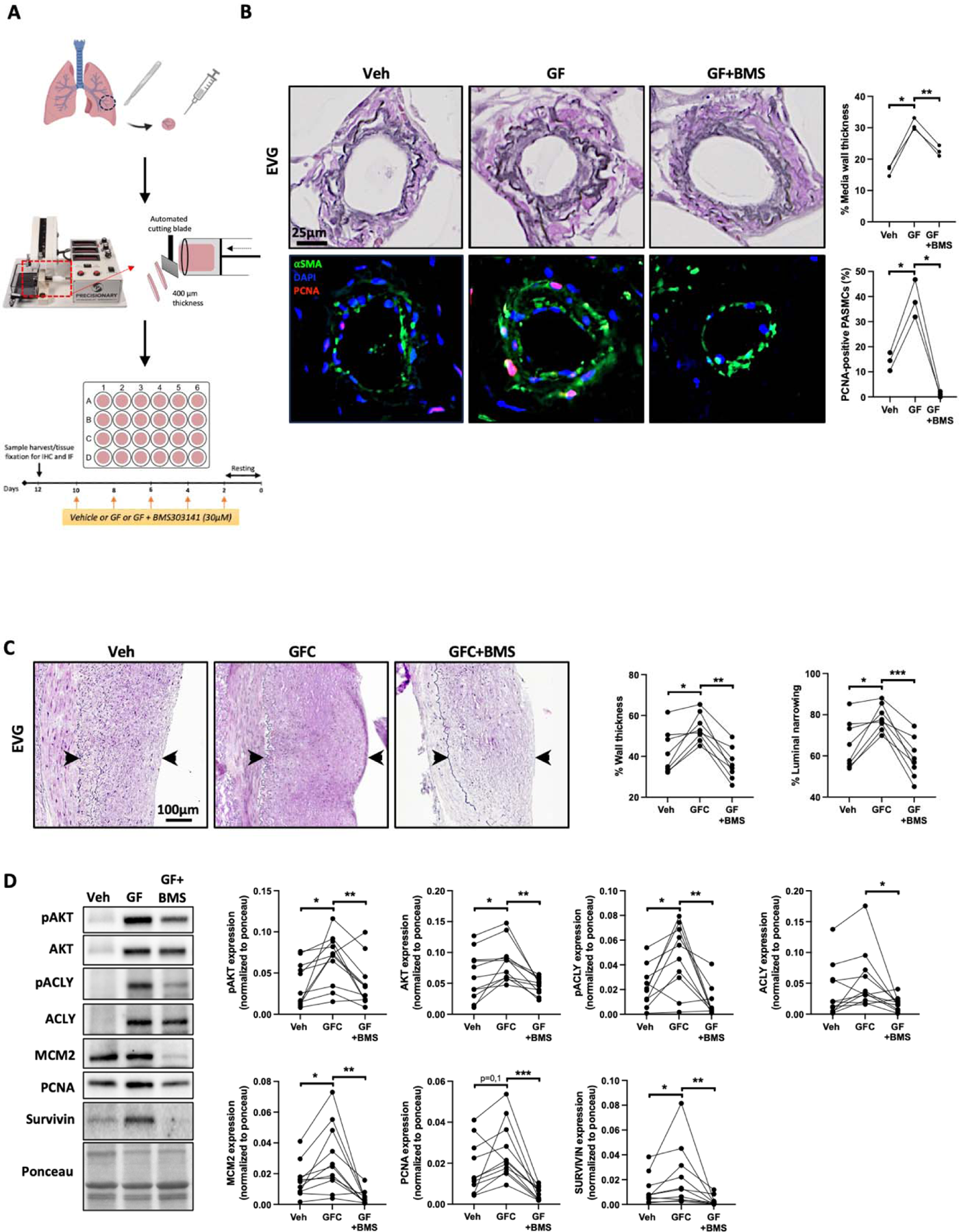
Pharmacological inhibition of ACLY reduces vascular remodeling in human precision-cut lung slices (PCLS) and cultured human coronary artery rings. **(A)** Experimental setup for PCLS. **(B)** Representative images of distal PAs stained with Elastica van Gieson (EVG) or labelled with proliferating cell nuclear antigen (PCNA) in PCLS prepared from control patients after exposure or not to a growth factor cocktail (GFC, FGF2+PDGF-BB+ET1) in presence or not of BMS-303141 for 10 days. Quantifications of medial wall thickness and PCNA-positive PASMCs (labelled with αSMA) in PCLS exposed or not to a GFC in presence or not of BMS-303141 are shown (n=3, *p<0.05, **p<0.01 repeated measured one-way ANOVA followed by Tukey’s post hoc analysis data represent mean ± SEM). Scale bar, 25μm. **(C)** Representative images of distal coronary rings stained with EVG after exposure or not to a GFC in presence or not to BMS-303141 for 5 days. The quantifications of wall thickness and luminal narrowing are shown. (n=8, *p<0.05, ***p<0.001 repeated measured one-way ANOVA followed by Tukey’s post hoc analysis data represent mean ± SEM). Scale bar, 100μm. **(D)** Representative Western blots and corresponding quantifications of pAKT, AKT, pACLY, ACLY, MCM2, PCNA and Survivin in coronary rings exposed or not to a GFC in presence or not to BMS-303141 for 5 days. (n=10 to 11, *p<0.05, **p<0.01, ***p<0.001 (repeated measured one-way ANOVA followed by Tukey’s post hoc analysis or Friedman followed by Dunn’s post hoc analysis).

To substantiate our findings, mechanically injured human coronary artery segments were exposed to growth factors in presence or not of ACLY inhibitor. Under stimulation of a growth factor cocktail (PDGF-BB, FGF2 and 7-ketocholesterol), rings of coronary artery isolated from patients demonstrated increased thickness of arterial wall and concomitant reduction of lumen area, as illustrated by histomorphometric analyses, and immunoblots for proliferative/survival markers (Fig. 8, C and D). Co-exposition with BMS-303141 resulted in less constrictive vessel remodeling and reduction of the expression of proliferative and survival markers (Fig. 8, C and D). Similarly, vessel rings obtained from human saphenous veins from patients undergoing coronary bypass surgery and incubated in presence of the growth factor cocktail for 5 days developed an inward remodeling (Fig. S16A), which was accompanied by an increased expression of cell proliferation and survival markers (Fig. S16B). Co-treatment with BMS repressed the remodeling process (Fig. S16B).

Together, these results provide additional compelling evidence that targeting ACLY may have therapeutic potential to prevent pathological vascular remodeling.

## DISCUSSION

The acquisition and maintenance of a synthetic phenotype by SMCs is a hallmark and driving process of arterial remodeling diseases such as PAH and systemic vascular stenosis. In this regard, continuous efforts have been made to delineate the mechanisms and factors involved in SMC phenotypic modulation. Nevertheless, our understanding remains incomplete, limiting the development of novel therapeutic approaches. In the present study, we demonstrated that increased activation of ACLY is used to initiate and support arterial remodeling in these two pathological contexts. Using *in vitro*, *ex vivo* and *in vivo* models, we provided converging evidence that interfering with ACLY inhibits SMC survival and proliferation, leading to attenuation of injury-induced constrictive remodeling and luminal narrowing. Specifically, we showed that, by regulating glucose and lipid metabolism, along with the pro-proliferative transcription factor FOXM1 and histone acetylation, ACLY supports multiple aspects responsible for the development of vascular lesions. Therefore, ACLY represents a novel and attractive therapeutic target for vascular remodeling diseases. Furthermore, our findings align with the concept that altered metabolism has a profound impact on chromatin remodeling and transcription, influencing cell function more broadly (*39, 40*).

While the overexpression of ACLY in vascular diseases has not been previously documented, our findings align with existing literature. Indeed, ACLY is known to be regulated by SREBPs transcription factors, which are activated by various mediators, including resistin (*41–43*). Intriguingly, the resistin/SREBPs axis is recognized for its involvement in CAD and PAH (*41–43*), providing additional support for the observed upregulation of ACLY in our study.

We demonstrated that inhibiting ACLY in both PAH and CAD SMCs results in a significant reduction in proliferation, migration, and apoptosis resistance as previously reported in cancer (*8*). These effects are associated with a decrease in the Warburg effect, *de novo* lipid synthesis (including cholesterol synthesis), and histone acetylation. Mechanistically, these effects are associated with a reduction in nuclear acetyl-CoA levels, accompanied by the downregulation of genes crucial for lipid synthesis and cell proliferation. Our computational investigation identified FOXM1 as a transcription factor responsible for the downregulation of genes in both PAH and CAD after ACLY inhibition. Experimental confirmation highlighted the upregulation of FOXM1 in both conditions, its repression by ACLY inhibition, and the mimicry of ACLY inhibition effects by specific FOXM1 inhibition. These findings not only confirm the importance of FOXM1 in vascular diseases(*22, 44, 45*) but also provide a novel regulatory mechanism of its activity under the control of ACLY, which could benefit many other pathologies.

Exploring the mechanisms by which ACLY regulates gene expression, we found strong evidence that the histone acetyltransferase GCN5 likely mediates the link between ACLY and histone acetylation. Inhibiting GCN5 activity mimicked the effects observed with ACLY inhibitors consistent with the notion that nuclear ACLY serves as a localized acetyl-CoA source for GCN5 activity(*46*) which in turn regulates FOXM1 expression through acetylation of the histones. However, it is important to acknowledge certain limitations in our study. Identification of the ACLY-regulated genes, particularly those directly regulated by GCN5 and FOXM1, remains unknown, necessitating further studies. Unraveling the specific genes under the direct influence of these key regulators will enhance our understanding of the intricated molecular pathways involved in vascular remodeling. For example, FOXM1 upregulation by ACLY-GCN5 axis may results from a direct acetylation of the histones within promoter region of FOXM1 allowing the binding of the transcription factors known to directly regulate its expression. Among all the identified transcription factors known to directly regulate FOXM1 expression (*47*) only FOXM1 itself is downregulated by ACLY in both PAH and CAD further highlighting the existence of a positive auto-regulation loop of FOXM1 in VSMC (*22*).

In addition to impact histone acetylation, the reduction in intracellular acetyl-CoA levels following ACLY inhibition is expected to influence the acetylation of non-histone proteins, contributing to alterations in their stability, cellular localization, and activity. This highlights how ACLY inhibition’s benefits are likely to be caused by a variety of molecular mechanisms that are difficult to rank. In connection with this, our data support the view that ACLY-mediated regulation of FOXM1 transcriptional activity results in part through the inhibitory acetylation of FOXO1 resulting in an overactivation of FOXM1 as previously reported (*48, 49*).

While our study demonstrates the effectiveness of specific SMC ACLY loss-of-function in preventing and reversing vascular remodeling in animal models of PAH and restenosis, potential benefits of pharmacological ACLY inhibition in other cell types, such as macrophages, cannot be excluded. Indeed, previous reports show that AKT phosphorylation, a key player in vascular diseases, regulates ACLY activity, and inhibition of ACLY activity prevents atherosclerosis lesions by downregulating plasma and tissue lipid elevations and suppressing inflammation. Further research is required to elucidate the specific role of ACLY in inflammatory cells and its contribution to PAH and neointimal hyperplasia lesions.

The strength of our study lies in the use of multiple models and methods to validate the role of ACLY. Consistency across patient-derived tissues, cultured SMCs, and animal models underscores the reproducibility and robustness of our findings.

In conclusion, the present study contributes to the standpoint that ACLY is a promising and multifaceted target for improving vascular lesions seen in PAH and CAD. These findings open avenues for future research, including detailed mechanistic studies and the development of ACLY inhibitors with high specificity and favorable pharmacokinetic profiles suitable for clinical trials.

## MATERIALS AND METHODS

### Experimental design

This study aimed to scrutinize the role of ACLY in the pathogenesis of vascular remodeling in PAH and CAD while investigating the potential therapeutic benefits of ACLY inhibition. This objective was addressed by (i) delineating the function of ACLY in driving vascular smooth muscle cell (VSMC) proliferation, (ii) elucidating the mechanisms through which ACLY mediates pathogenic metabolism and epigenetic activity, orchestrating VSMC reprogramming in CAD and PAH, and (iii) evaluating therapeutic effects through ACLY inhibition and knockdown in systemic and PAH rodent models.

Cell culture experiments were conducted at least three times, with each replicate performed in triplicate. The number of animals per group was calculated to detect a minimum 20% difference between experimental and control groups, ensuring 80% power and a 10% standard deviation. The selection of unique patient samples was primarily determined by clinical availability. Animals of the same sex, genotype, and similar body weight were generated and randomly assigned to different experimental groups, with no exclusions from analyses.

Investigators conducting hemodynamic data collection and histologic analysis were blinded to the groups. All animal experiments received approval from the Laval University ethics committee CPAUL2 (CPAUL, #2019-311, #2020-616). Ethical approval for human tissue use adhered to the standards of the Declaration of Helsinki, with all experimental procedures approved by institutional review boards at the CRIUCPQ ethics committee (CER#20773, CER # 22197, CER #20841). Informed consent for this study conformed to the highest ethical standards.

### Human Patient Samples

Informed consent was diligently secured for all tissue sampling procedures. For samples derived from World Symposium on Pulmonary Hypertension (WSPH) Group 1 PAH, diagnosis was meticulously established by an expert physician utilizing specific criteria: elevated mean pulmonary arterial pressure (mPAP) exceeding 20 mmHg, pulmonary capillary wedge pressure below 15 mmHg, and pulmonary vascular resistance exceeding 3 Wood units, determined through right heart catheterization. In PAH cases, the expert physician adjudicated the diagnosis by systematically excluding left heart disease, hypoxic lung disease, or chronic thromboembolic disease.

Non-PAH human lung specimens utilized for precision-cut lung slices (PCLS) were sourced from three distinct categories: 1) excessively large donor lungs, 2) warm autopsy samples obtained from donors without any lung diseases, or 3) healthy lung sections (verified by a pathologist) derived from individuals undergoing lung surgery for tumor removal.

For tissue samples obtained from CAD patients, saphenous veins were acquired during Coronary Artery Bypass Grafting (CABG). Coronary arteries from CAD patients were sourced from individuals undergoing heart transplantation. CAD diagnosis was meticulously confirmed by an expert physician based on criteria involving the presence of stent and/or CABG. Vascular smooth muscle cells (VSMCs) were meticulously isolated from remodeled coronary arteries dissected by expert pathologists from the Institut Universitaire de Cardiologie et de Pneumologie de Québec.

### Statistical Analysis

Numerical quantifications from in vitro experiments using cultured cells, as well as in situ quantifications of transcript expression and physiological experiments using rodents or human reagents, are presented as mean ± SEM. Immunoblot images are representative of experiments repeated at least three times and quantified using [specific method/standard]. Micrographs are representative of experiments within each relevant cohort.

The normality of the data distribution was assessed through the Shapiro-Wilk test. To compare means between two sample groups with normal data distribution, an unpaired Student’s t-test was employed, while non-normally distributed data underwent Mann-Whitney testing. In the context of comparisons among different groups, one-or two-way analysis of variance (ANOVA) tests were conducted, and for normally distributed data, subsequent Tukey’s or Bonferroni’s post hoc analyses were performed. Conversely, Kruskal-Wallis tests, followed by Dunn’s post hoc analysis, were utilized for non-normally distributed data when comparing different groups. In the case of paired data, multi-repeated ANOVA was applied, with Tukey’s post hoc analysis for normally distributed data and Friedman’s test followed by Dunn’s post hoc analysis for non-normally distributed data. A significance level of P < 0.05 was adopted to establish statistical significance. All graphs and statistical analyses were completed using GraphPad Prism (ver.9.5.1).

## Acknowledgements

We thank the IUCPQ Biobank of the Quebec Respiratory Health Research Network as well as the department of cytology and pathology from the IUCPQ for providing access to tissue and clinical data.

## Author contributions

Conceptualization: OB, SB, SP, PV, CR, YG

Methodology: CR, YG, SEL, MS, RK, SM, MM, SBB, AB, CT, NF, ED, JP, FP Investigation: CR, YG, SEL, MS, RK, SM, MM, SBB, AB, CT, NF, ED, JP, FP

Visualization: OB, SB, SP, PV, CR, YG Funding acquisition: OB, SB

Project administration: Supervision: SP, OB, SB

Writing – original draft: YG, SP, OB, SB

## Data and materials availability

All data are available in the main text or the supplementary materials.

