## supplemental materials and figures for "ATP Citrate Lyase Drives Vascular Remodeling Diseases Development Through Metabolic-Epigenetic Reprograming"

**List of Supplementary Materials**

**Materials and methods**

**Human tissues and cells**

Experimental procedures using human tissues or cells were performed in conformity with the principles outlined in the Declaration of Helsinki and were approved by the Laval University and the Institut Universitaire de Cardiologie et de Pneumologie de Québec (IUCPQ) Biosafety and Ethics Committees (CER#20773, CER # 22197, CER #20841). Tissues were obtained from patients who had previously given signed informed consent. The patients’ characteristics are described in Supplemental Tables S2 and S3

The pulmonary arteries smaller than 1 mm in outer diameter were isolated from both PAH and Non-PAH patients and PASMCs enzymatically isolated as previously described*(1)*. Coronary arteries from both non-CAD and CAD patients were obtained from explanted heart at the time of surgery. Briefly, patients with stent and/or bypass graft were considered as CAD patients while patients with no clinical history of atherosclerosis were considered has non-CAD patients. CoASMCs were enzymatically isolated as previously described*(1)*.

**Ex-vivo human coronary artery and saphenous vein culture:**

Coronary arteries from explanted heart and saphenous veins (obtained from patients undergoing ischemic heart disease) were isolated at the time of the surgery and cut in rings of 1 to 2cm long. All the rings received mechanical endothelial denudation using clamp and were placed in culture for 5 days with (PDGF-BB 30ng/mL+FGF2 30ng/mL; 7-keto cholesterol 10µM) in presence or not to ACLY inhibitor BMS-303141 at 30µM. The medium and treatments were changed every 24h. After 5 days rings, were harvested for histological and western blotting-based experiments.

**Ex-vivo human pulmonary cut lung slices (PLCS):**

PCLS were obtained from tumor-free lung explants collected during lung resection surgeries performed on patients with cancer. Lung tissues were processed maximum within 1-2 days after resection. Briefly, lung was warmed and inflated delicately with warm (37°C) low melting agarose (3%) in HBSS/HEPES solution (10mM, pH7-7,5). After the agarose solidified, tissue was cut in 2cm x 1cm x 1cm rectangle. Before precision cutting with the compresstome vibratome (VF-310-OZ), the tissue samples were embedded with agarose solution. The cutting process was executed using the following settings: a cutting frequency of 3 Hz, a cutting speed of 3 mm/s, and slices with a thickness of 400 μm. Each slice was sequentially placed in one well of a 24-well plate and then in an incubator set a temperature of 37°C, providing 5% of CO2. Lung sections were cleaned during the first two hours with agitation and regular media change. Viability assay was immediately performed with MTT and LDH assays (Sigma M2128) according to the manufacturer’s instruction. At day 2, lung sections were cultured in DMEM supplemented with 1% FBS, 0.1% antibiotic-antimycotic solution and 0,01% gentamycin and stimulated or not with a cocktail of growth factor to induce vascular remodeling (PDGF-BB 30ng/mL, FGF-2 30ng/mL, endotheline-1 100ng/ml) in combination with BMS-303141(30μM) or its vehicle. Media was changed every 48 hours for a treatment duration of 10 days. Slices were fixed in 4% FA, embedded in paraffin, and processed for histological measurements.

**Animals and *in vivo* treatments**

All animal studies were approved by the Animal Ethics Committee of Université Laval (CPAUL, #2019-311, #2020-616) and in accordance with the guidelines of the Canadian Council on Animal Care. The animals were kept under standard laboratory with free access to tap water and food.

Mice with *loxP* sites flanking exon 9 of the *Acly* gene (*Acly*^flox/flox^, *Acly^tm1.1Welk^*/Mmjax, The Jackson Laboratory)*(2)* were crossed with *Tagln-Cre* mice (B6.Cg-Tg(Tagln-cre)1Her/J, The Jackson Laboratory) or *Myh11-CreER^T2^* mice (B6.FVB-Tg[Myh11-cre/ERT2]1Soff/J, The Jackson Laboratory) expressing tamoxifen-inducible CRE under the SMC-specific Myh11 promoter, to generate *Acly*^+/flox^;Tg^+/Tagln-Cre^, *Acly*^flox/flox^;Tg^+/Tagln-Cre^, and *Acly*^flox/flox^;Tg^+/Myh11-CreERT2^ mice. Mice were genotyped by PCR using the genotyping protocols suggested by the supplier. In the first part of the study, the Sugen/Hypoxia (Su/Hx) rat model of PAH was used. Briefly, after acclimatization adult male Sprague-Dawley rats (Charles River laboratories; 200-250 in body weight) received a single subcutaneous injection of the VEGF receptor antagonist SU5416 (20mg/kg, APExBIO) and were then maintained in hypoxia (10% O_2_) for 21 days. Then, these rats returned to normoxia for 2 additional weeks. Su/Hx rats with established PAH (3 weeks post-SU5416 injection) were randomly assigned to either a vehicle control group (0.5% Carboxymethyl cellulose and 0.025% Tween 20), a group that received BMS-303141 (5mg/kg, once daily by oral gavage) for 2 weeks. Control rats were maintained in normoxic conditions. Regarding PH induction in mice, adult wildtype (*Acly*^+/flox^ and *Acly*^flox/flox^), *Acly*^+/flox^;*Tg*^+/Tagln-Cre^, *Acly*^flox/flox^;*Tg*^+/Tagln-Cre^, and *Acly*^flox/flox^;Tg^+/Myh11-CreERT2^ mutant mice (8-11 weeks of age; weight, 17-30g) were weakly injected SU5416 at 20mg/kg once a week for 3 weeks. After the first injection of Sugen, mice were exposed to hypoxia for 3 weeks. Adult male mice were then injected with Tamoxifen (2mg/mouse intraperitoneal), once a day for 5 consecutive days for two weeks in normoxia condition.

In the second part of the study, animal models of carotid injury were used. The left common carotid artery of the anesthetized Sprague Dawley rats (aged 6-8 weeks, 350-450g) was exposed through a midline cervical incision, and blood flow to the site was temporarily interrupted by clamps. A small incision was made, and a flexible metal guide wire (2 mm in diameter) was introduced into the artery and passed up and down 5-10 times with rotation through the clamped section to denude the endothelium. After mechanical denudation, the arteriotomy was closed with monofilament Ethilon non-absorbable 9-0 suture and the blood flow was reestablished by removing the clamps. Only male rats were used for this study to avoid the interference of hormones. Injured rats were randomly assigned to received either BMS-303141 (5mg/kg) or its vehicle (0.5% Carboxymethyl cellulose and 0.025% Tween 20) via oral gavage, once a day beginning the day of wire injury for two weeks. Rats in the sham operation group received midline cervical incision and left carotid artery isolation without endothelial denudation. To induce neointima formation in the mouse carotid artery*,* a carotid ligation model was used *(3)*. Briefly, the left common carotid arteries of 8-10-week-old WT and mutant mice were exposed and ligated with PROLENE monofilament non-absorbable 6-0 suture. Note that for experiments conducted with *Acly*^flox/flox^;Tg^+/Myh11-CreERT2^ mice, only males were used as the transgene is inserted into the Y chromosome *(4)*.

In accordance with the 3Rs principals (Replacement, Reduction and Refinement) for better ethical use of animals in research, saphenous veins obtained from dogs subjected to coronary artery bypass graft (CABG) surgery were reused to assess the relationship between saphenous veins remodeling and ACLY expression. Briefly, as in humans 30% of the saphenous veins CABG in dogs failed due to intimal hyperplasia. Tissues were harvested 45 days post-surgery and intimal hyperplasia along with ACLY expression were quantified as described in the manuscript.

**Echocardiographic and hemodynamic measurements in PAH models**

At the end of treatment, animals were initially anesthetized with 3%–4% isoflurane and maintained with 2% during procedures. Transthoracic echocardiography was performed in mice using a Vevo 3100 (Visualsonics) imaging system. Pulmonary hemodynamics were assessed through images acquired from a parasternal short-axis and long-axis views, as previously described *(5)* and according to current guidelines *(6)*. Next, hemodynamic parameters, including RV systolic pressure (RVSP) and mean PA pressure (mPAP) were measured blindly by closed chest right heart catheterization (RHC) (SciSence catheters), as previously described *(5)*. After hemodynamics had been recorded, the animals were killed, and their tissues were harvested.

**Histological and immunofluorescence analysis**

Tissues were fixed with 4% paraformaldehyde, paraffin-embedded and serially sectioned at 5µm. Tissue sections were deparaffinized with xylene and rehydrated through a graded ethanol series and processed for Elastica van Gieson (EVG, #Sigma 1159740002) or immunofluorescence staining as described below. For immunofluorescence staining, tissue sections were immerged in 10mM sodium citrate buffer (pH 6.0) and heated in a microwaveable pressure cooker for 15min. Following antigen retrieval, the sections were blocked with normal goat serum (10%) in PBS for 2h at room temperature. Then, sections were incubated with indicated primary antibodies (Table S4) in a humidified chamber at 4°C overnight. After washes in PBS, sections were incubated for 1 h at room temperature with appropriate fluorescent-dye conjugated secondary antibodies (Table S4). Sections were mounted onto coverslips using DAPI (4′,6-diamidino-2-phenylindol) Fluoromount G mounting medium. Sections were viewed and examined by microscopy using a digital slice scanner (Axio Scan Z1, Zeiss) or an Axio Observer microscope (Zeiss), and images were acquired using Zen system (Zeiss).

**Morphometric assessment of pulmonary vascular remodeling and neointimal formation**

In PAH animal models, the degree of PA medial wall thickness was measured and expressed as follows: Percent of wall thickness = [(external diameter–internal diameter)/external diameter] × 100. At least 15 randomly selected distal PA per animal were measured in a blinded fashion to assess vascular remodeling and proliferation. For CAD models, the extent of neointima formation was evaluated by quantifying the luminal, intimal (internal elastic lamina (IEL) − Lumen), and medial (external elastic lamina (EEL) − IEL) areas, as previously described *(7)*. The intima-to-media (I/M) ratio was calculated from the mean of these determinations. The luminal obliteration was defined as the percentage of area within the IEL blocked by the neointima.

**Cell culture and treatments**

PASMCs and CoASMCs were grown in smooth muscle cell basal medium supplemented with smooth muscle cell growth kit (Cell Applications) and maintained at 37°C in a 5% CO_2_ air humidified incubator. Cells were used at passages 4 to 9 for experiments (Table S2 and S3). BMS-303141 (ACLY pharmacological inhibitor) and Butyrolactone 3 (BA3, a GCN5 pharmacological inhibitor) were purchased from MedChemExpress and Cedarlane, respectively. The compounds were dissolved in DMSO and were applied to the PASMCs and CoASMCs at various concentrations for the indicated time. siACLY (#L-004915-00-0005, Dharmacon), siFOXM1 (#sc-270048, Santa Cruz Biotechnology) and negative control siRNA (#4390846, Thermo Fisher Scientific) were transfected at a final concentration of 10nM and 50nM, respectively, with Lipofectamine RNAiMAX (Thermo Fisher Scientific) according to the manufacturer’s instructions.

### **In vitro proliferation, apoptosis, and migration measurements**

To assess cell proliferation and apoptosis, PASMCs and CoASMCs were cultured basal medium designed for the culture of smooth muscle cells (#310-500, Cell Applications) supplemented with cell growth supplement, antibiotics, and FBS (#311-GS, Cell Applications). After treatments (48 post exposure to drug or 72h post siRNA transfection), cells were fixed with 4% formaldehyde diluted in phosphate-buffered saline (PBS) 1X for 15 min at room temperature, washed with PBS 1X and then permeabilized for 15 min in 0.2% Triton X-100 in PBS. Cell proliferation was determined with Ki67 labeling. Apoptosis was evaluated by Annexin V assay, as previously described*(8, 9)*. The Ki67 proliferative and Annexin V apoptotic index were calculated by counting the number of positive-staining cells divided by the total number of DAPI-positive cells multiplied by 100. For each cell line, experiments were performed in triplicate and at least 500 cells per condition were counted. A wound healing assay was used to assess the migration ability of PAH-PASMCs and CAD-CoASMCs following knockdown of ACLY. Briefly, a total of 2.5×10^5^ transfected cells were seeded into each well of six-well plates and cultured until a monolayer of cells had formed. A 200 *μ*L pipette tip was used to create similar size of vertical and horizontal scratches in the cell layer for each group. Next, scratched cells were removed by gently rinsing PBS twice; and the remaining cells were continually cultured in serum-free medium for a 24h period. The wound closure was monitored using a microscope and images of the same region of interest were taken at selected time points. Images were analyzed with ImageJ software and Wound Healing Size Tool plugin *(10)*. Since BMS treatment requires a 48-hour duration, the migration assay was not performed with BMS to circumvent the need for administering an additional drug dose after the 24-hour pretreatment period.

**RNA Sequencing and analysis**

RNA was isolated from cells using the Direct-zol RNA MicroPrep Kit (Zymo Research) according to the manufacturer’s instructions. DNase I (RNeasy MinElute Cleanup kit; QIAGEN) was used to remove genomic DNA contamination, while RNA integrity and quantity was assessed using a NanoDrop spectrophotometer (Thermo-Fisher). RNA sequencing was performed by the CHU de Québec-Université Laval Research Center genomics platform. Briefly, samples were processed using NEBNext Ultra II directional RNA library prep kit (New England Biolabs), with library preparations sequenced on the Novaseq 6000 Sequencing System (Illumina) with 100 base-pair single end reads, resulting in approximatively 30 million single end reads per sample. FASTQ files from the Novaseq instrument were examined for sequencing quality and adapters contamination using MultiQC v1.13 *(11)*. Reads with low quality were removed prior to further analyses and adapters sequences were removed from the reads using Trimmomatic v0.36 *(12)*. Alignment of trimmed reads to the human genome (hg38) was performed using the STAR aligner software v2.7.10 *(13)* and the data were converted to gene‐level expression using RSEM v1.1.17*(14)*. Gene counts from the datasets were then used to assess differential expression using DESEQ2 v1.39.3 *(15)*. High-throughput sequencing data used in this study have been deposited in NCBI’s Gene Expression Omnibus and are accessible through GEO Series accession number ***GSE to be determined upon acceptation*** and ***GSE to be determined upon acceptation***. An absolute fold change of ≥1.5 and adjusted p-values ≤0.05 were used to define genes whose expression was significantly altered. Kyoto Encyclopedia of Genes and Genomes (KEGG) pathway enrichment analysis was performed using ShinyGO (version 0.77) *(16)*.

**Quantitative Real Time PCR**

Total RNA from tissues or cells was isolated using TRIzol reagent (Invitrogen), according to the manufacturer’s instructions. One µg of RNA was then converted into cDNA using the qScript Flex cDNA Synthesis Kit (Quanta Bio). Quantitative PCR was performed in triplicate on the QuantStudio 7 Flex real-time PCR system (Applied Biosystems). Amplification of a single DNA product was confirmed by melting curve analysis and gel electrophoresis. The primers used are presented in Supplemental Table S5. The relative expression level of each target was determined by the ΔΔCt method. Transcript levels of genes of interest were normalized to 18S mRNA.

**Western blotting**

Tissues or cells were lysed in RIPA buffer supplemented with protease and phosphatase inhibitor cocktail, and the protein concentrations were measured by Bradford assay. Equal amount of proteins were resolved by sodium dodecyl sulfate–polyacrylamide gel electrophoresis (SDS-PAGE), and then transferred to a nitrocellulose or polyvinylidene difluoride membrane. The membranes were then blocked with either 5% BSA or 5% non-fat dry milk in TBS-T buffer, and incubated with primary antibodies overnight at 4°C. Then the membranes were rinsed 3 times with TBS-T buffer, 10 minutes each, and incubated with appropriate horseradish peroxidase (HRP)-conjugated secondary antibodies diluted in blocking buffer for 2 hour at RT. Finally, antibodies were revealed using ECL reagents (Perkin–Elmer) and labeled proteins were detected with the imaging Chemidoc MP system (Bio-Rad Laboratories). Protein expression was quantified using the Image lab software (Bio-Rad Laboratories) and normalized to ponceau or amido black as indicated. The sources and working dilutions or primary and secondary antibodies are listed in Table S4.

**Cell fractionation**

Subcellular fractionation to isolate nuclear and cytoplasmic fractions was done using a nuclear extraction kit (#ab219177, Abcam), as previously described*(17)*. Protein concentration was measured by Bradford assay. Separation between the nuclear and cytosolic fractions was verified by blotting for the cytosolic protein α/β Tubulin and the nuclear protein Histone H3.

**LC-MS/MS analyses**

Incubations were stopped by removing the cell supernatants, which were immediately mixed 0.5 volume of cold (4˚ Celcius) methanol. Cell pellets were scraped in ice-cold PBS and immediately denatured with 0.5 volume of cold methanol. Methanol levels then were adjusted to 50%. Deuterated internal standards were added and samples were processed as previously described *(18)*. LC-MS/MS analyses were next performed using a Shimadzu 8050 triple quadrupole mass spectrometer using the documented gradient and multiple reaction monitoring transitions previously described *(19, 20)*

**Acetyl-CoA measurement**

The acetyl-CoA content was determined on nuclear fractions using the PicoProbe Acetyl CoA Assay Kit (fluorometric) per manufacture instructions (#ab87546, Abcam).

**Histone extraction**

This procedure utilized the histone extraction kit (# ab113476, Abcam).

**Mitochondrial oxygen consumption rate (OCR) and extracellular acidification rate (ECAR) assay**

Oxygen consumption rate (OCR) and basal extracellular acidification rate (ECAR) of PAH-PASMCs and CAD-CoASMCs were measured using the Seahorse XF24 Bioanalyser (Seahorse Bioscience). Forty eight hours after siRNA transfection, PAH-PASMCs or CAD-CoASMCs were seeded in Seahorse 24-well tissue culture plates at a density of 3.5-4 × 10^4^ cells/well and allowed to adhere for 24 h. Prior to the mitostress assay, cell confluence was confirmed, the media was changed to Agilent Seahorse XF RPMI medium pH 7.4 supplemented with 25 mM glucose, 4 mM L-glutamine and 1 mM sodium pyruvate, and the cells were equilibrated for 1 h at 37 °C in a non-CO_2_ incubator. OCR was measured under basal conditions. Uncoupled and maximal OCR were determined after the addition of 8μM oligomycin and 45μM fluoro-carbonyl cyanide phenylhydrazone (FCCP). Rotenone (10nM) was used to inhibit mitochondrial respiration. Prior to the glycostress assay, cell confluence was confirmed, the media was changed to Agilent Seahorse XF RPMI medium pH 7.4 supplemented with 2 mM L-glutamine and the cells were equilibrated for 1 h at 37 °C in a non-CO_2_ incubator. ECAR was measured after the addition of 80 mM glucose. Maximal ECAR was determined after the addition of 9µM oligomycin. 2-deoxy-gluocse (2-DG, 500mM) was used to inhibit glycolysis. OCR and ECAR values were normalized to the total protein concentration per well assessed by a Bradford protein assay (Bio-Rad Laboratories) after completion of the XF assay.

**Oil red O staining (ORO)**

CoASMCs and PASMCs were plated on coverslip into twenty-four well plate. After treatments media of cells was removed and rinsed 2 times with PBS and cells were fixed with 4% FA for 15 minutes. FA was discarded and cells were washed two times with PBS. ORO was performed as previously described *(21)*. Then, cells were washed with distillated water and mounted with DAPI Fluoromount G medium. Cells were observed and pictured by microscopy using an Axio Observer microscope (Zeiss), and images were acquired using Zen system (Zeiss) by fluorescence with Differential Interference Contrast. Images were analyzed with ImageJ software (NIH).

**Filipine III staining**

Filipin III staining kit (Sigma, SAE0087) was used to measure free cholesterol in vitro. Briefly, after treatment, cells were rinsed three times with PBS and fixed with 3% FA for 1h at room temperature (RT). FA was discarded and celles were washed 3 times with PBS. Cells were then incubated during 10 min with 1ml of 1,5mg glycine/ml PBS. Next, cells were stained with 1ml of filipin working solution (0,05mg/ml in PBS with 10%FBS) for 2h at Room temperature. Finally, cells were rinsed 3 times with PBS then mounted with Fluoromount G medium (without DAPI). Cells were observed and pictured by microscopy using an Axio Observer microscope (Zeiss), and images were acquired using Zen system (Zeiss) by fluorescence.

**Fig S1 to S16**

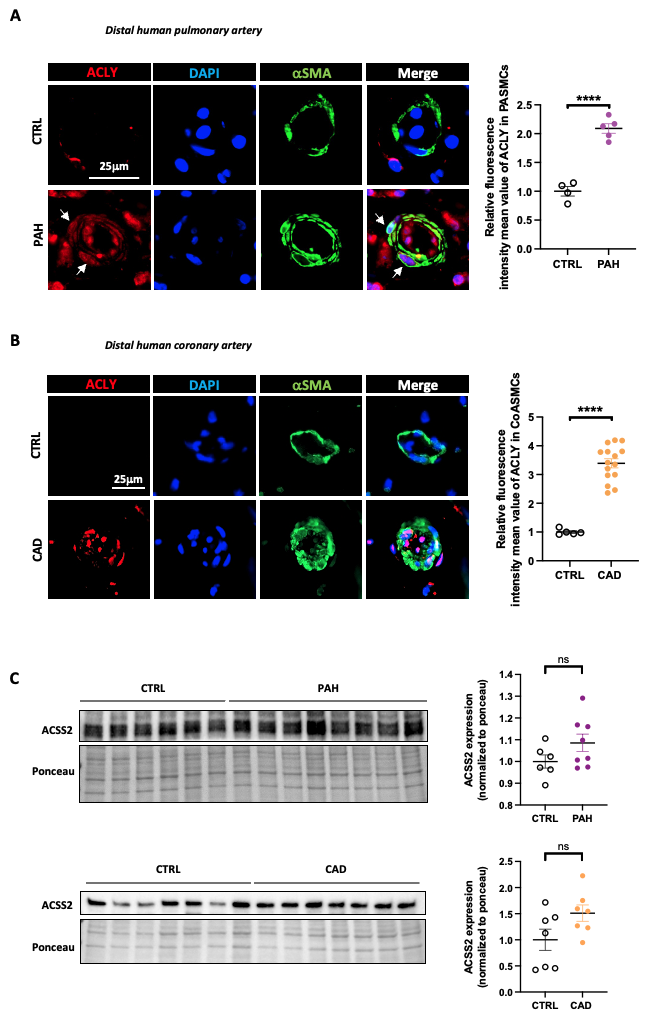

**Fig. S1. Increased expression of ACLY in pulmonary artery and coronary artery smooth muscle cells from PAH and CAD patients, respectively**

**(A)** Representative immunofluorescence images of ACLY (red) and αSMA (green) in distal pulmonary arteries (PAs) from healthy donors and patients with PAH. This staining was completed with DAPI (blue) nuclear counterstaining for cell identification and numbering within tissues. The immunofluorescent quantification for ACLY in PASMCs is shown (n=4 or 5; ****p<0.0001, unpaired Student’s *t* test; data represent mean ± SEM). Scale bars, 25 μm. **(B)** Representative immunofluorescence images of ACLY (red) and αSMA (green) in coronary arteries from healthy donors and patients with CAD. This staining was completed with DAPI (blue) nuclear counterstaining for cell identification and numbering within tissues. The immunofluorescent quantification for ACLY in SMCs is shown (n=5 or 15; ****p<0.0001 unpaired Student’s *t* test; data represent mean ± SEM). Scale bars, 25 μm. **(C)** Western blots and corresponding quantifications of ACSS2 in PASMCs and CoASMCs isolated from healthy donors and patients with PAH or CAD. (n=6 to 8; unpaired Student’s *t* test; data represent mean ± SEM).

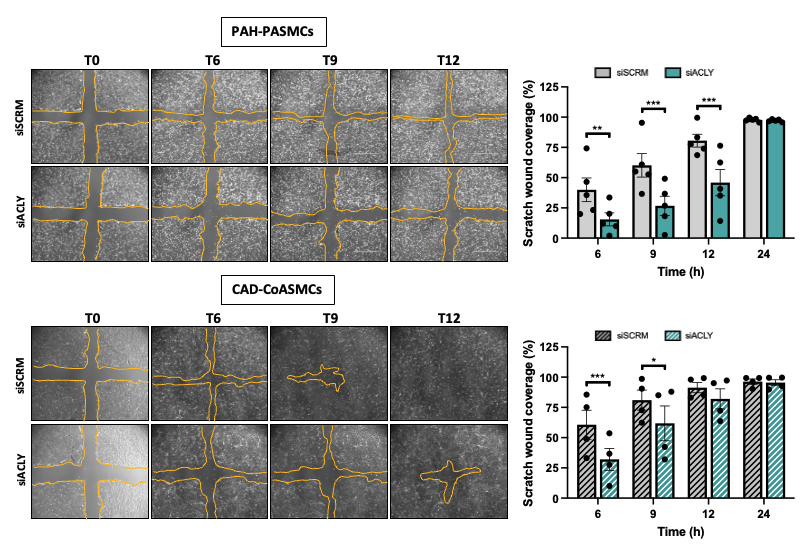

**Fig. S2. ACLY promotes migration of PAH-PASMCs and CAD-CoASMCs**

Representative bright-field images of wound-healing assay for confluent monolayers of PAH-PASMCs and CAD-CoASMCs subjected or not to ACLY knockdown at different time points. Right: bar graphs of percentage of wound closure at 6, 9, 12, and 24h post-injury. (n=4 or 5; *p<0.05, **p<0.01, ***p<0.001, 2-way ANOVA followed by Bonferroni’s post hoc analysis; data represent mean ± SEM). Scale bar, 1000 μm.

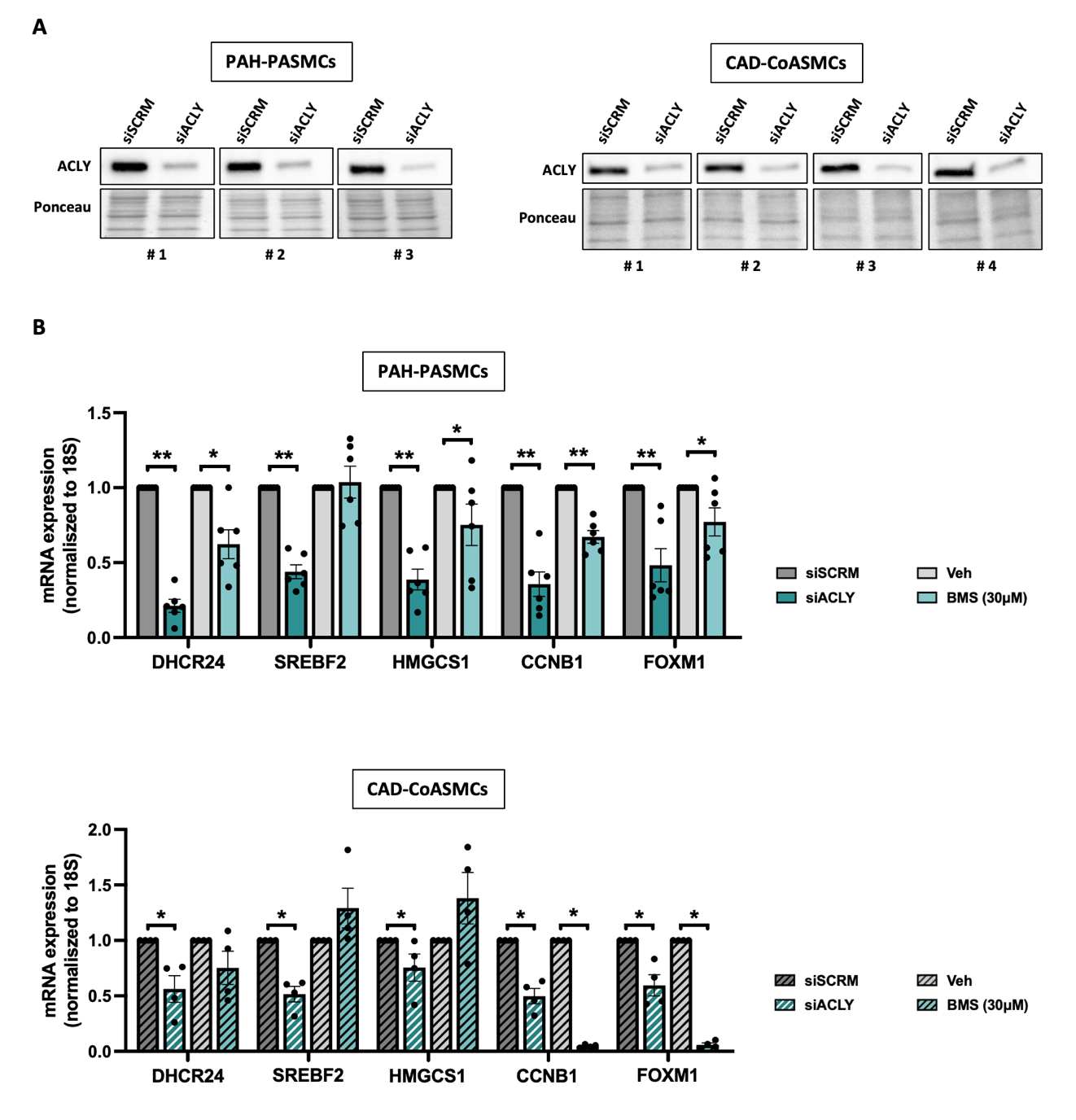

**Fig. S3. Validation of transcriptomic results**

**(A)** Western blots of ACLY in PAH-PASMCs and CAD-CoASMCs subjected or not to ACLY knockdown for 72h using siRNA. **(B)** Relative mRNA expression of selected genes in PAH-PASMCs and CAD-CoASMCs subjected or not to ACLY inhibition using siACLY or BMS-303141 (30μM) for 72h and 48h, respectively. (n=4 or 6; *p<0.05, **p<0.01, Mann-Whitney’s test; data represent mean ± SEM).

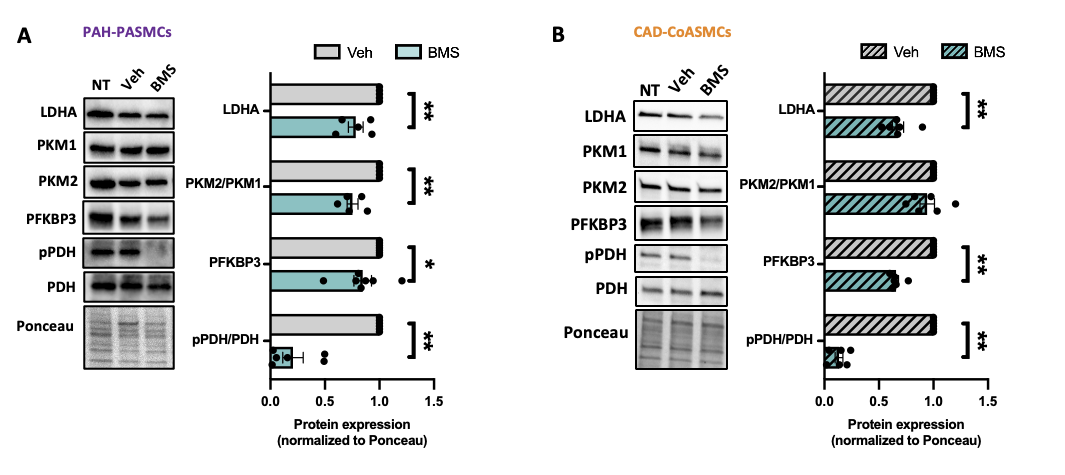

**Fig. S4. Impact of ACLY inhibition on expression of glycolysis markers in PAH-PASMCs and CAD-CoASMCs**

**(A** and **B)** Representative Western blots, and corresponding quantifications of LDHA, PKM1, PKM2, PFKBP3, phospho-PDH (pPDH), and PDH expression in PAH-PASMCs (A) and CAD-CoASMCs (B) subjected or not to ACLY inhibition using BMS-303141 (30mM) for 48h. (n=5 or 7; *p<0.05, **p<0.01 Mann-Whitney’s test data represent mean ± SEM).

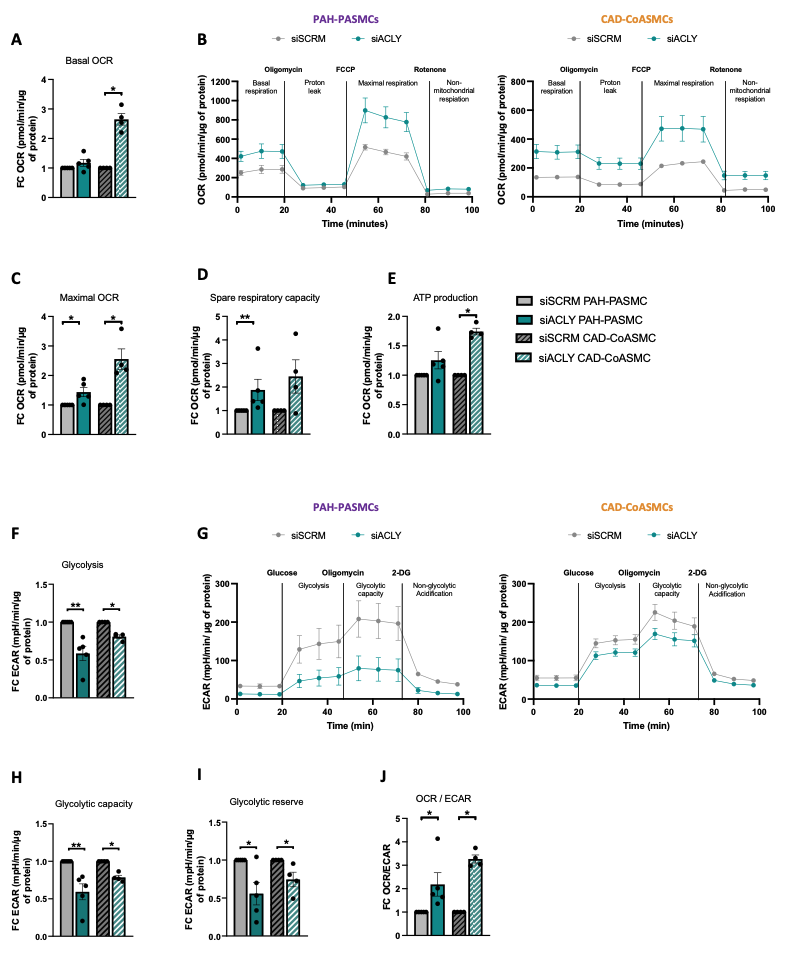

**Fig. S5. Impact of ACLY inhibition on PAH-PASMCs and CAD-CoASMCs bioenergetics.**

**(A)** Basal oxygen consumption rate (OCR) measurement by Seahorse in PAH-PASMCs and CAD-CoASMCs subjected or nor to ACLY knockdown for 72h (n=4 or 5; *p<0.05, **p<0.01; Mann-Whitney’s test data represent mean ± SEM). **(B)** Representative curves for Seahorse measurements of OCR in PAH-PASMCs and CAD-CoASMCs transfected with either a siRNA that targets ACLY or a scrambled siRNA for 72h. **(C** to **E)** Quantifications of maximal OCR (C), spare respiratory capacity (D), ATP production (E), and basal extracellular acidification rate (ECAR) (F) in PAH-PASMCs and CAD-CoASMCs subjected or not to ACLY knockdown for 72h (n=4 or 5; *p<0.05, **p<0.01 Mann-Whitney’s test; data represent mean ± SEM). **(G)** Representative curves for Seahorse measurements of ECAR in PAH-PASMCs and CAD-CoASMCs transfected with either a siRNA that targets ACLY or a scrambled siRNA for 72h. **(H** to **J)** Quantifications of glycolytic capacity (H), glycolytic reserve (I), and OCR/ECAR ratio (J) in PAH-PASMCs and CAD-CoASMCs subjected or not to ACLY knockdown for 72h (n=4 or 5; *p<0.05, **p<0.01 Mann-Whitney’s test.; data represent mean ± SEM).

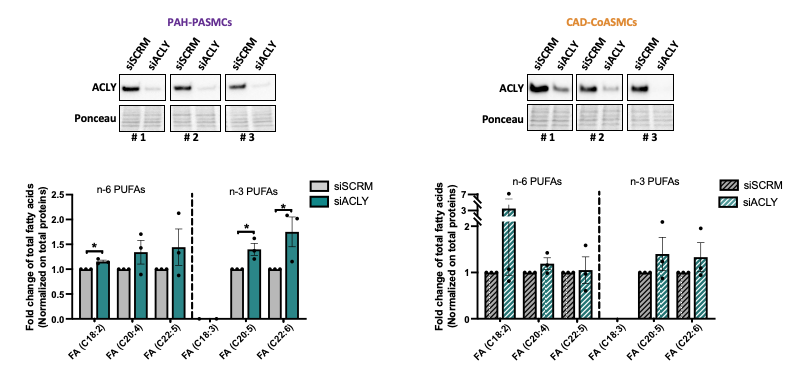

**Fig. S6. Impact of molecular ACLY inhibition on levels of polyunsaturated fatty acids in PAH-PASMCs and CAD-CoASMCs**

**(A)** Representative Western Blots of ACLY following siACLY exposure for 72h on both CAD-CoASMCs and PAH-PASMCs. **(B)** PUFAs measurements using mass spectrometry in the same cells (n=3 *p<0.05, Mann-Whitney’s test; data represent mean ± SEM).

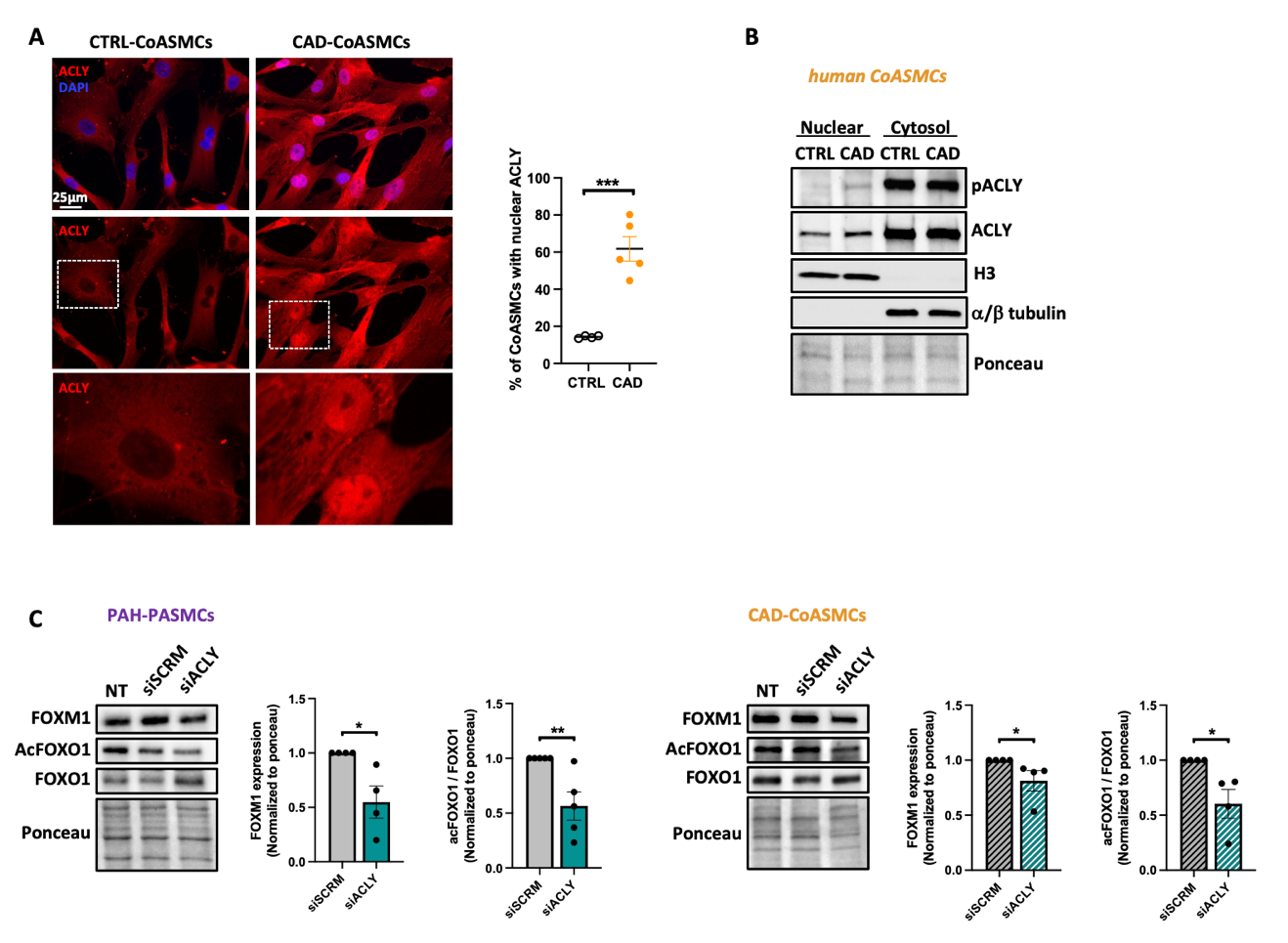

**Fig. S7. Subcellular localization of ACLY in CoASMCs from control and CAD patients and impact of its inhibition on FOXM1 and FOXO1**

**(A**) Representative fluorescent images of control and CAD-CoASMCs labeled with ACLY. Scale bar, 25μm. **(B)** Representative Western blots (of 3 independent experiments) for phospho-ACLY (S455), ACLY, α/β tubulin, and histone H3 (H3) in nuclear and cytosolic extracts from control and PAH-PASMCs. **(C)** Representative Western blots and corresponding quantification of FOXM1, acetyl-FOXO1 (K294) and FOXO1 in PAH-PASMCs and CAD-CoASMCs subjected or not to ACLY inhibition using siRNA for 72h (n=4-5 *p<0.05, **p<0.01 Mann-Whitney’s test; data represent mean ± SEM).

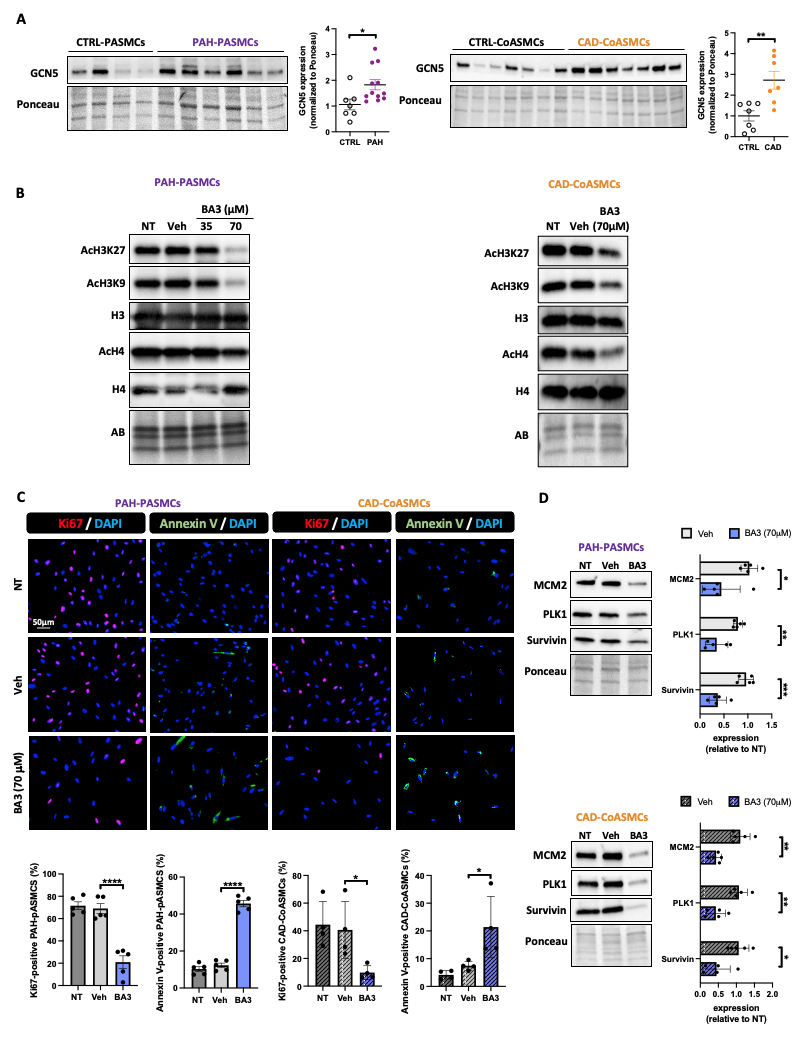

**Fig. S8. Impact of GCN5 inhibition on PAH-PASMCs and CAD-CoASMCs**

**(A)** Representative Western blots, and corresponding quantifications of GCN5 in PASMCs and CoASMCs isolated from control and patients with PAH (Mann-Whitney’s test) or CAD (unpaired Student’s *t* test) (n= 7 or 12; *p<0.05, **p<0.01, data represent mean ± SEM). **(B)** Representative Western blots, confirming efficacy of GCN5 inhibition on AcH3K27, AcH3K9, H3, AcH4 and H4 in PAH-PASMCs and CAD-CoASMCs treated or not with butyrolactone 3 (BA3) for 48h. **(C)** Representative fluorescent images, and corresponding quantifications of Ki67-labeled (red) and Annexin V-labeled (green) PAH-PASMCs and CAD-CoASMCs treated or not with BA3 (70μM) for 48h or exposed or not to BMS-303141 (30μM) for 48h (n=4 or 5; *p<0.05, ****p<0.001, one-way ANOVA followed by Tukey’s post hoc analysis; data represent mean ± SEM). **(D)** Representative Western blots, and corresponding quantifications of MCM2, PLK1, and Survivin in PAH-PASMCs and CAD-CoASMCs treated or not with butyrolactone 3 (BA3) for 48h (n= 5; *p<0.05, **p<0.01, ***p<0.001 unpaired Student’s *t* test); data represent mean ± SEM).

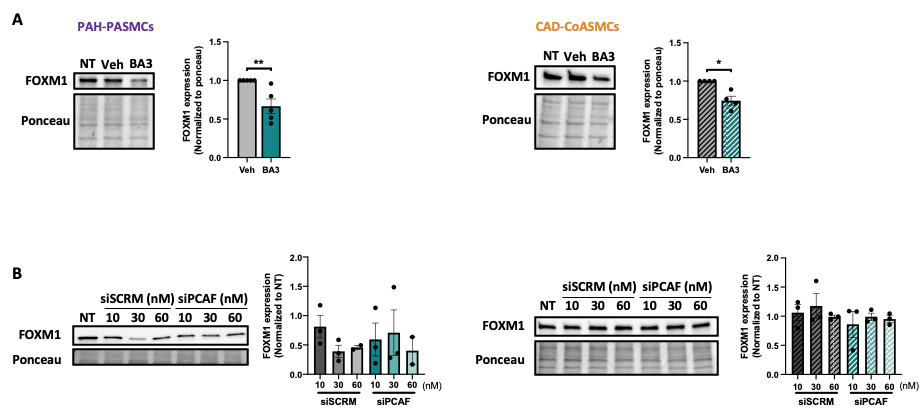

**Fig. S9. Impact of GCN5 inhibition on FOXM1 expression in PAH-PASMCs and CAD-CoASMCs**

**(A)** Representative Western blots, and corresponding quantifications of FOXM1 in PAH-PASMCs and CAD-CoASMCs treated with BA3 (70µM) or its vehicle (DMSO) for 48h (n= 4 or 5; *p<0.05, **p<0.01 Mann-Whitney’s test; data represent mean ± SEM). **(B)** Representative Western blots and corresponding quantifications of FOXM1 in PAH-PASMCs and CAD-CoASMCs subjected or not to PCAF knockdown using siRNA for 72h (n= 3; Mann-Whitney’s test; data represent mean ± SEM).

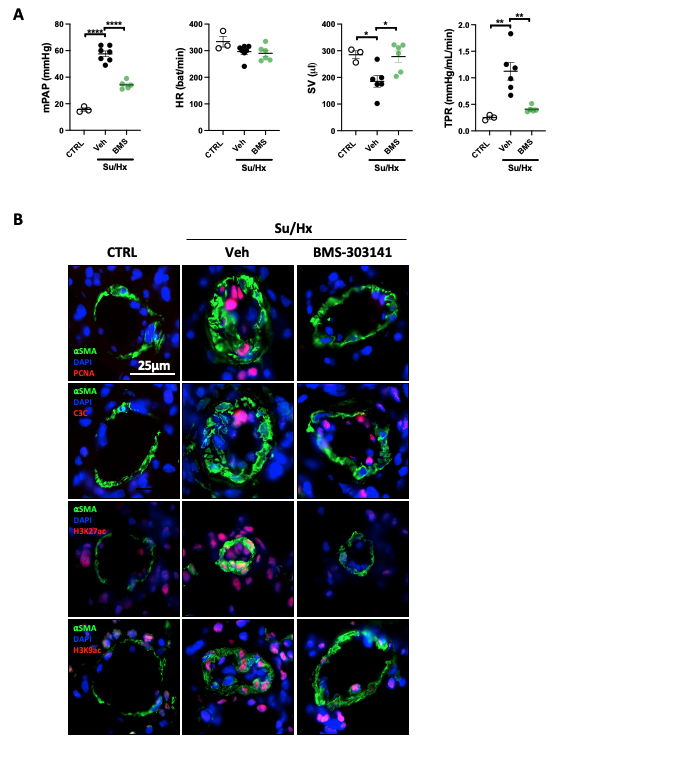

**Fig. S10. Pharmacological inhibition of ACLY improves established PAH in Su/Hx rats**

**(A)** mean pulmonary artery pressure (mPAP), heart rate (HR), stroke volume (SV), and total pulmonary resistance (TPR) in BMS-303141-treated Su/Hx PAH rats versus controls as assessed by right heart catheterization (n=3 to 7; *p<0.05, **p<0.01, ***p<0.001, one-way ANOVA followed by Tukey’s post hoc analysis; data represent mean ± SEM). **(B)** Representative immunofluorescence images of PCNA, cleaved caspase 3 (C3C), acH3K27, acH3K9, and SMA (green) in distal pulmonary arteries (PAs) from control, Su/Hx+Veh and Su/Hx+BMS-303141 rats. This staining was completed with DAPI (blue) nuclear counterstaining for cell identification. Scale bar, 25μm.

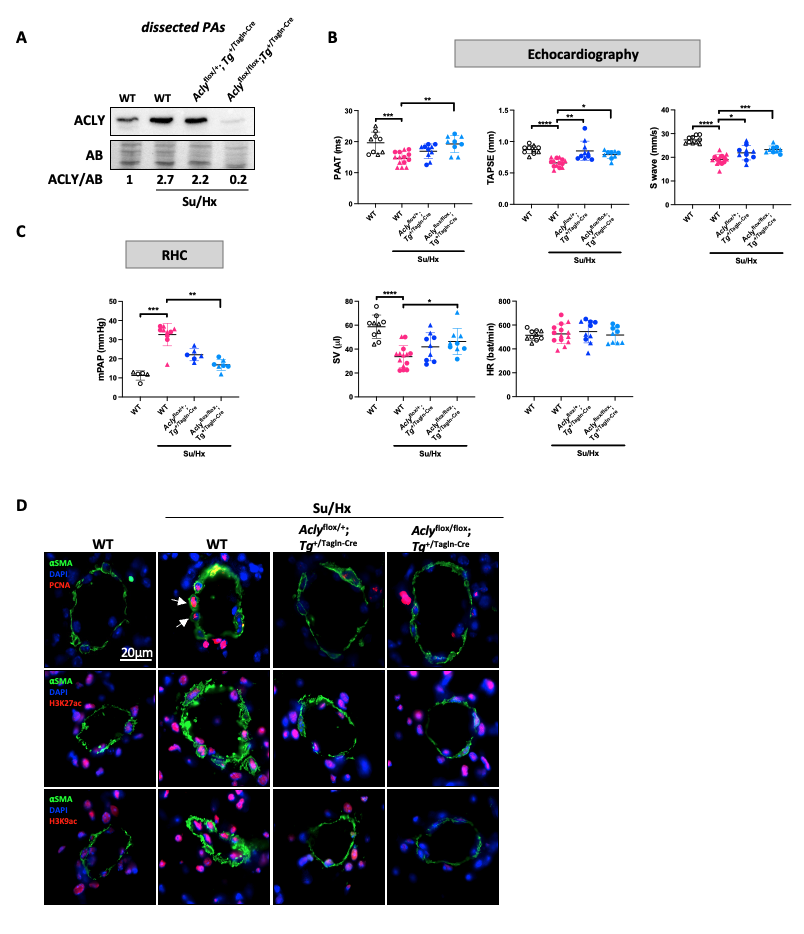

**Fig. S11. *Acly* loss-of-function targeted to smooth muscle cells confers protection against Su/Hx-induced PAH in mice**

**(A)** Western blot of ACLY in pooled PAs from wildtype (WT, Acly^flox/flox^), Su/Hx-exposed WT, *Acly*^+/flox^;Tg^+/Tagln-Cre^, and *Acly*^flox/flox^;Tg^+/Tagln-Cre^ mice. **(B)** Pulmonary artery acceleration time (PAAT) (one-way ANOVA followed by Tukey’s post hoc analysis), tricuspid annular plane systolic excursion (TAPSE) (Kruskal-Wallis followed by post hoc Dunn’s test), S wave (one-way ANOVA followed by Tukey’s post hoc analysis), stroke volume (SV) (one-way ANOVA followed by Tukey’s post hoc analysis) and heart rate (HR) (one-way ANOVA followed by Tukey’s post hoc analysis) determined by echocardiography at the end of the protocol in WT, *Acly*^+/flox^;Tg^+/Tagln-Cre^, and *Acly*^flox/flox^;Tg^+/Tagln-Cre^ mice exposed to either normoxia (Nx) or Sugen/Hypoxia (Su/Hx) for 3 weeks. Male and female mice are indicated by circles and triangles, respectively (n=9 to 14; *p<0.05, **p<0.01, ***p<0.001, ****p<0.0001; one-way ANOVA followed by Tukey’s post hoc analysis data represent mean ± SEM). **(C)** mean pulmonary artery pressure (mPAP) determined by right heart catheterization in control, Su/Hx-exposed WT, *Acly*^+/flox^;Tg^+/Tagln-Cre^, and *Acly*^flox/flox^;Tg^+/Tagln-Cre^ mice (n=5 to 10; ***p<0.001, ****p<0.0001, Kruskal-Wallis followed by post hoc Dunn’s test; data represent mean ± SEM). **(D)** Representative immunofluorescence images of distal pulmonary arteries (PAs) labeled for PCNA, acH3K27, acH3K9, and aSMA (green) in WT, *Acly*^+/flox^;Tg^+/Tagln-Cre^, and *Acly*^flox/flox^;Tg^+/Tagln-Cre^ mice exposed to either normoxia or Su/Hx for 3 weeks. Nuclei are counterstained with DAPI (blue). Scale bar, 20mm.

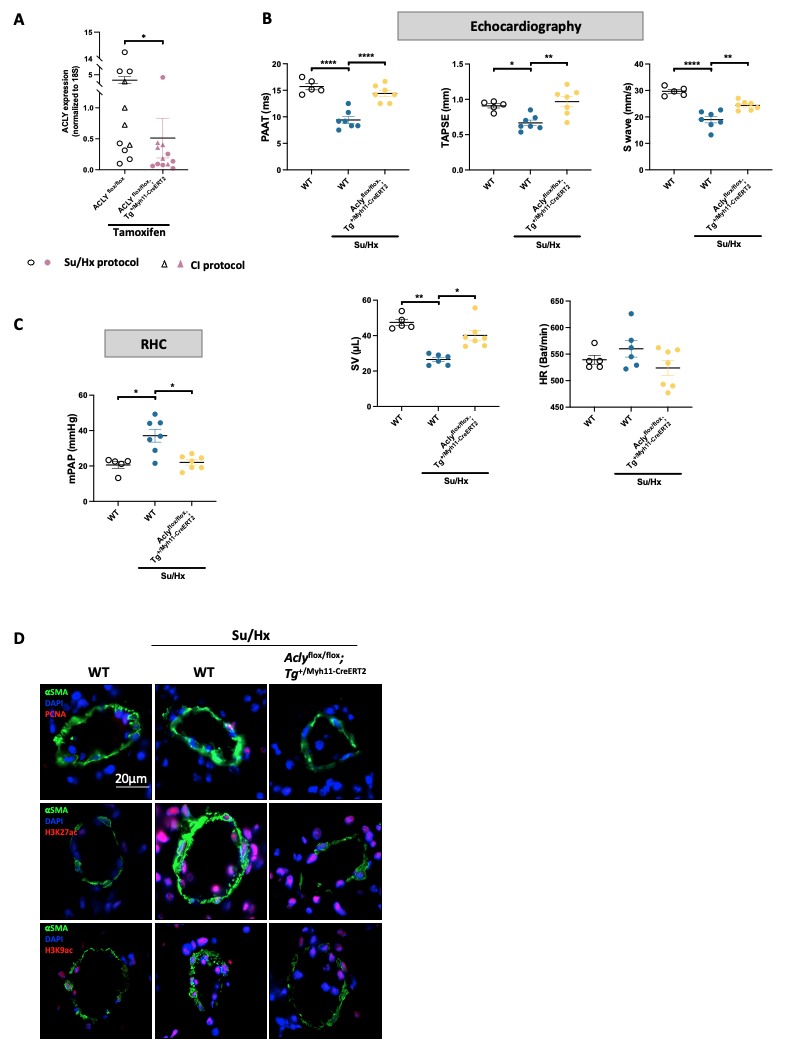

**Fig. S12. Inducible *Acly* loss-of-function targeted to smooth muscle mitigates PAH development in the Su/Hx mice**

**(A)** Relative mRNA expression of *Acly* in pooled aorta from tamoxifen-treated *Acly*^flox/flox^ (WT) and *Acly*^flox/flox^;Tg^+/Myh11-CreERT2^ male mice at the end of the Su/Hx and carotid ligation protocols. Mice used for Su/Hx and carotid ligation protocols are indicated by circles and triangles, respectively (n=11 to 12; *p<0.05, Mann-Whitney’s test; data represent mean ± SEM). **(B)** Pulmonary artery acceleration time (PAAT); Tricuspid annular plane systolic excursion (TAPSE), S wave (one-way ANOVA followed by Tukey’s post hoc analysis), stroke volume (SV) (Kruskal-Wallis followed by post hoc Dunn’s test), and heart rate (HR) (Kruskal-Wallis followed by post hoc Dunn’s test) determined by echocardiography at the end of the protocol in Tamoxifen-treated Su/Hx-exposed WT and *Acly*^flox/flox^;Tg^+/Myh11-CreERT2^ mice vs controls. (n=5 to 7; *p<0.05, **p<0.01, ****p<0.0001; data represent mean ± SEM). **(C)** mean pulmonary artery pressure (mPAP) determined by right heart catheterization in Tamoxifen-treated Su/Hx-exposed WT and *Acly*^flox/flox^; Tg^+/Myh11-CreERT2^ mice vs controls. (n=5 to 7; *p<0.05 Kruskal-Wallis followed by post hoc Dunn’s test; data represent mean ± SEM). **(D)** Representative immunofluorescence images of distal pulmonary arteries (PAs) labeled for PCNA, acH3K27, acH3K9, and αSMA (green) in Tamoxifen-treated WT and *Acly*^flox/flox^; Tg^+/Myh11-CreERT2^ mice exposed to either normoxia or Su/Hx for 3 weeks. Nuclei are counterstained with DAPI (blue). Scale bar, 20μm.

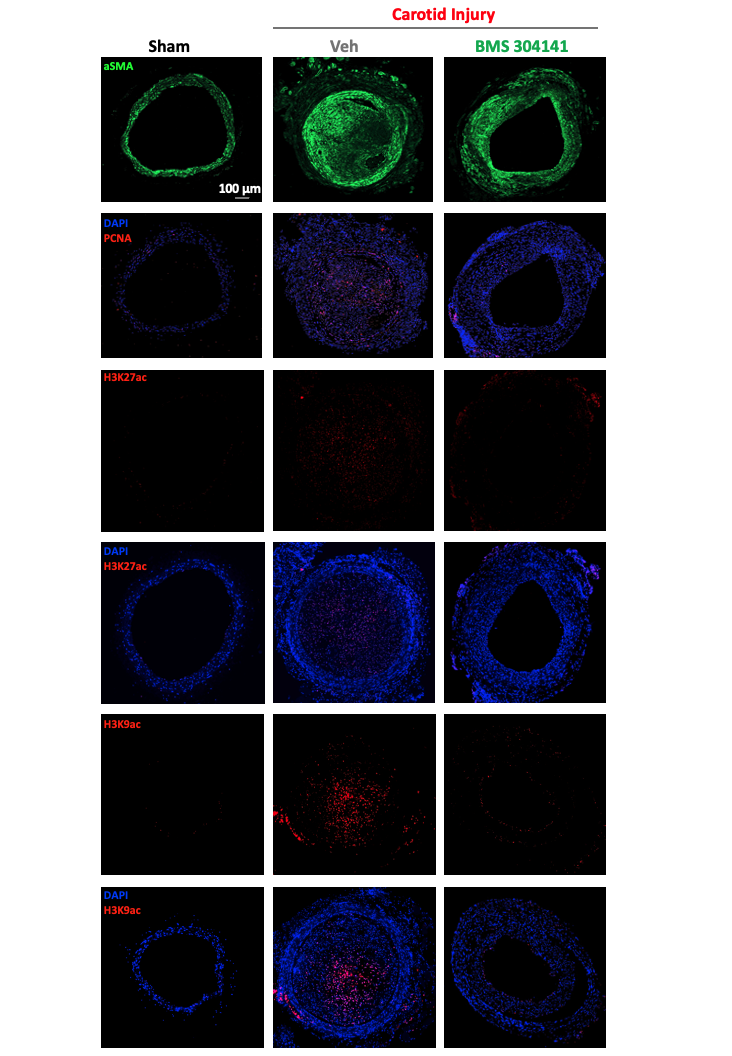

**Fig. S13. Pharmacological ACLY inhibition confers protection against carotid remodeling induced by carotid injury.**

Representative immunofluorescence images of left carotid arteries labeled for PCNA**,** acH3K27, acH3K9, and αSMA (green) in rats subjected or not to vascular injury and treated or not with BMS-303141 for 2 weeks. The staining was completed with DAPI (blue) nuclear counterstaining for cell identification. Scale bar, 100m.

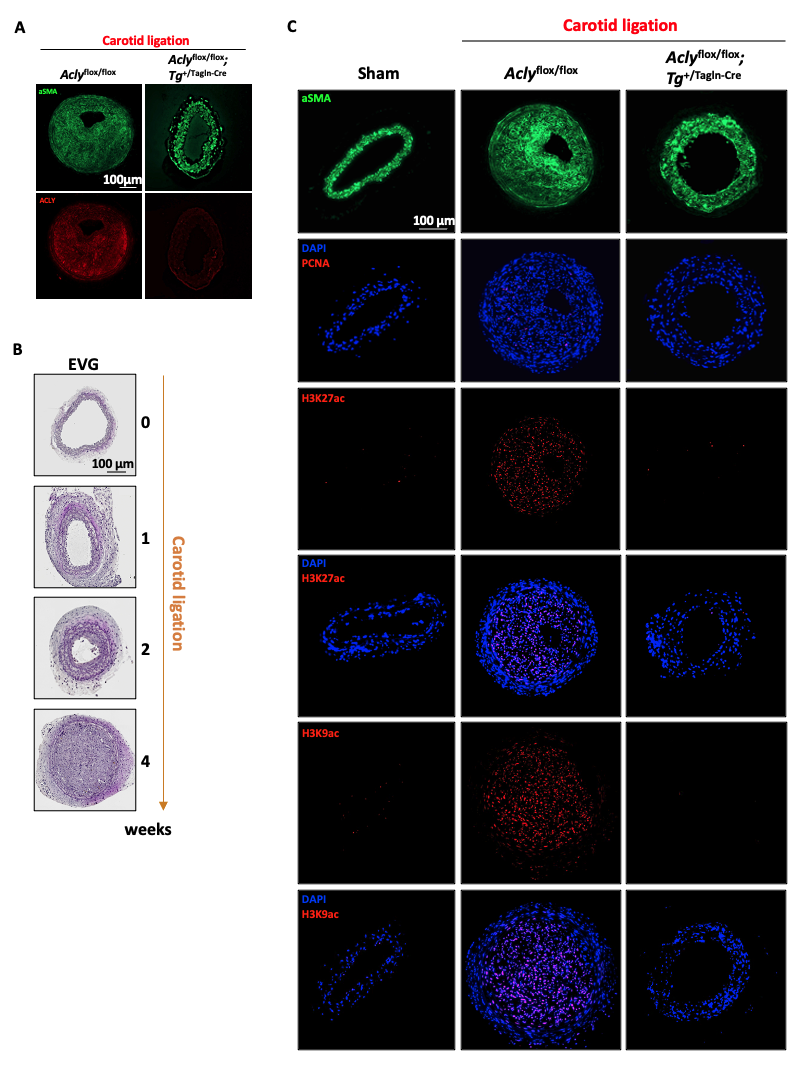

**Fig. S14. Role of ACLY in coronary artery remodeling**

(A) Representative immunofluorescence images of left carotid arteries labeled for ACLY (red) and αSMA (green) in *Acly*^flox/flox^ (WT) and *Acly*^flox/flox^; Tg^+/Tagln-Cre^ mice subjected to carotid ligation. Scale bar, 100m. **(B)** Representative photomicrographs of Elastica van Gieson (EVG)-stained left coronary arteries in mice at different time points after ligation. Scale bar, 100μm**. (C)** Representative immunofluorescence images of left carotid arteries labelled with PCNA (red), acH3K27 (red), acH3K9 (red), and αSMA (green) in WT and *Acly*^flox/flox^; Tg^+/Tagln-Cre^ mice subjected or not to carotid ligation. This staining was completed with DAPI (blue) nuclear counterstaining. Scale bar, 100μm

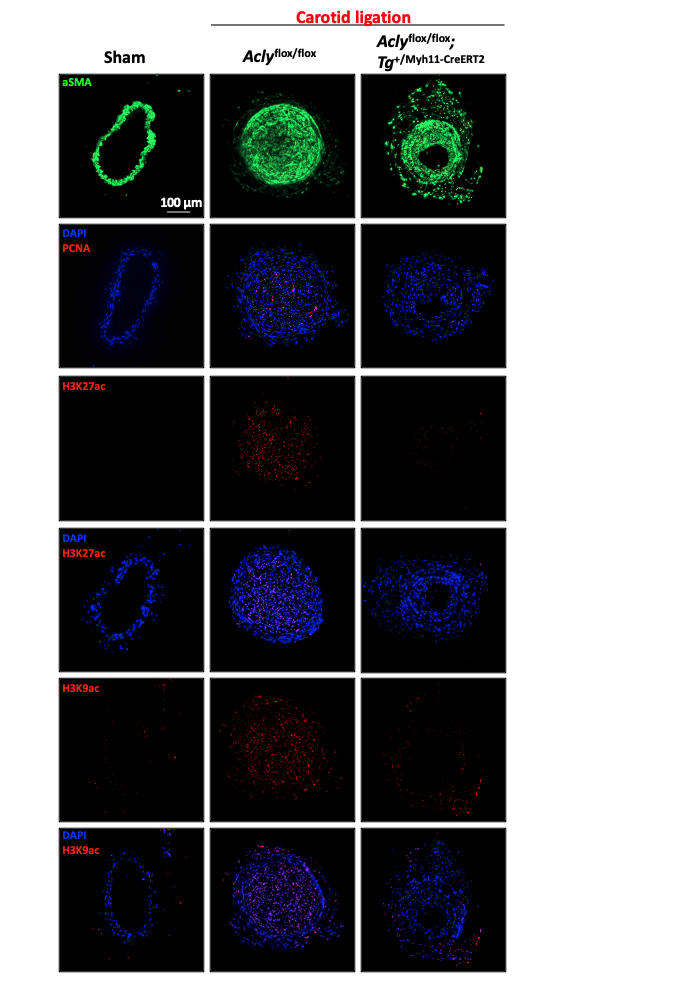

**Fig. S15. *Acly* loss-of-function targeted to smooth muscle cells confers protection against carotid remodeling induced by carotid ligation in mice**

Representative immunofluorescence images of left carotid arteries labelled with PCNA (red), acH3K27 (red), acH3K9 (red), and αSMA (green) in Tamoxifen-treated WT and *Acly*^flox/flox^;Tg^+/Tagln-Cre^ mice subjected or not to carotid ligation. This staining was completed with DAPI (blue) nuclear counterstaining. Scale bar, 100μm.

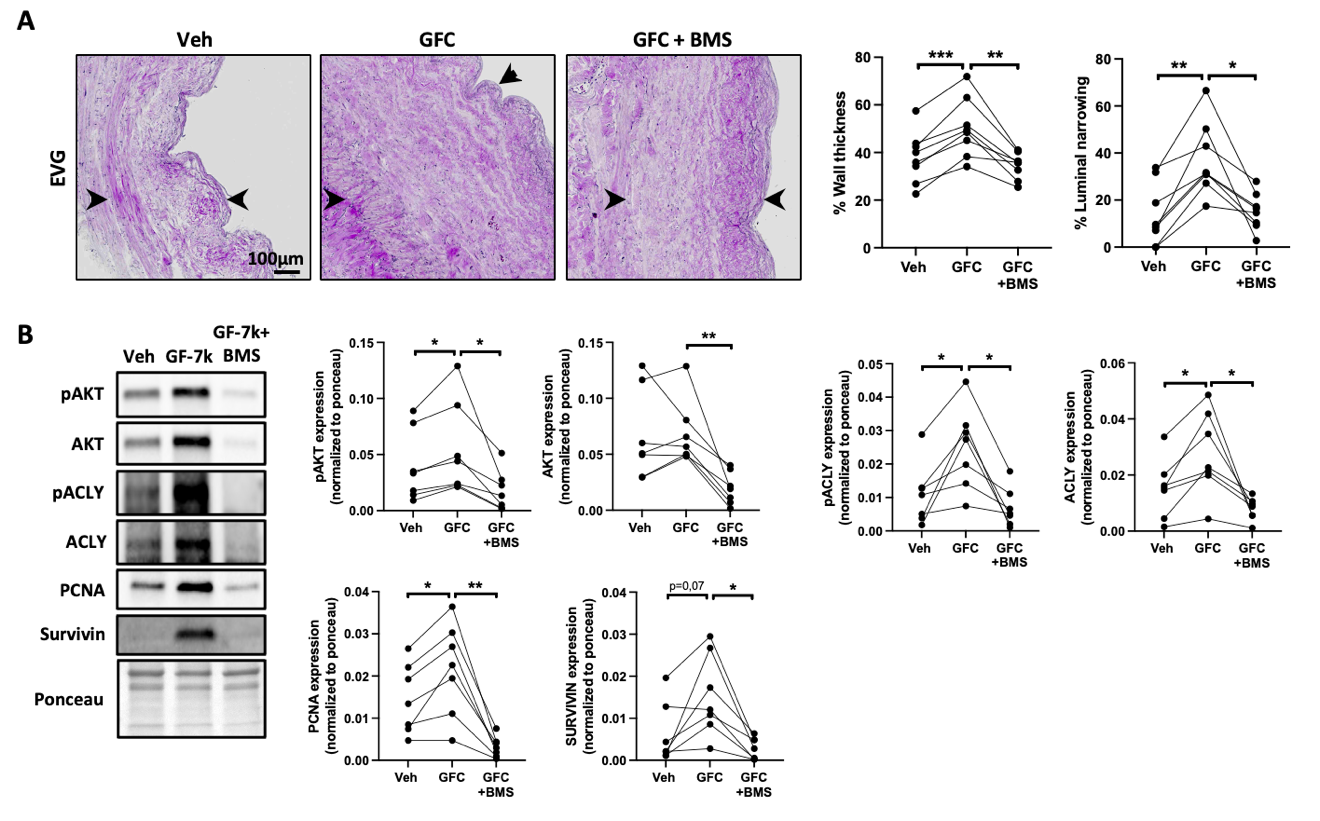

**Fig. S16. ACLY inhibition reduces vascular remodeling in cultured human saphenous veins rings**

**(A)** Representative images of human sephanous rings stained with Elastica van Gieson (EVG) after exposure or not to a growth factor cocktail (PDGF-BB; FGF2; 7-keto cholesterol) in presence or not to BMS-303141 for 5 days. The quantifications of wall thickness and luminal narrowing are shown. (n=8, *p<0.05, **p<0.01 repeated measured ANOVA followed by Tukey’s post hoc analysis). Scale bar, 100μm. **(B)** Representative Western blots, and corresponding quantifications of pACLY, ACLY, PCNA and Survivin in human saphenous veins rings exposed or not to a GFC in presence or not to BMS-303141 for 5 days. (n=7, *p<0.05, **p<0.01, ***p<0.001 repeated measured ANOVA followed by Tukey’s post hoc analysis).

**Table S1 to S4**

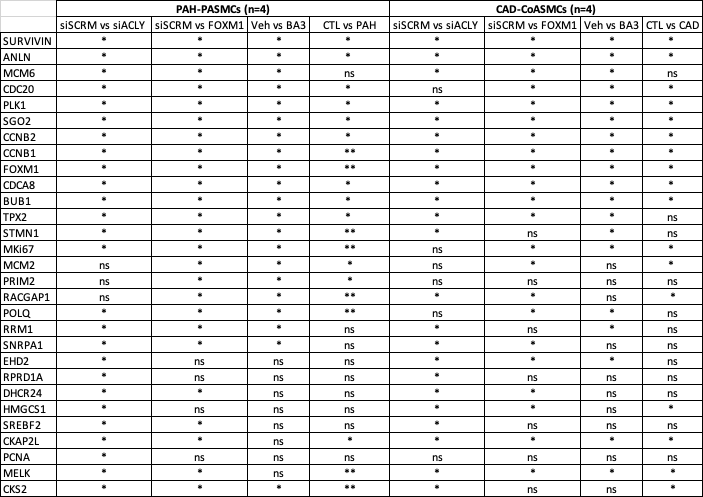

**Table S1. qRT-PCR results**

Genes expression differences measured by qRT-PCR in PASMCs and CoASMCs isolated from 8 control (4 PASMCs and 4 CoASMCs) and 4 PAH-PASMCs and 4 CAD-CoASMCs exposed to either scrambled siRNAs, siACLY, siFOXM1 (72h) or vehicle (DMSO 0.1%) or BA3 (GCN5 inhibitor) 48h at 70uM. *p<0,05 and ** p<0,01 unpaired student t-test or Mann-Whitney test.

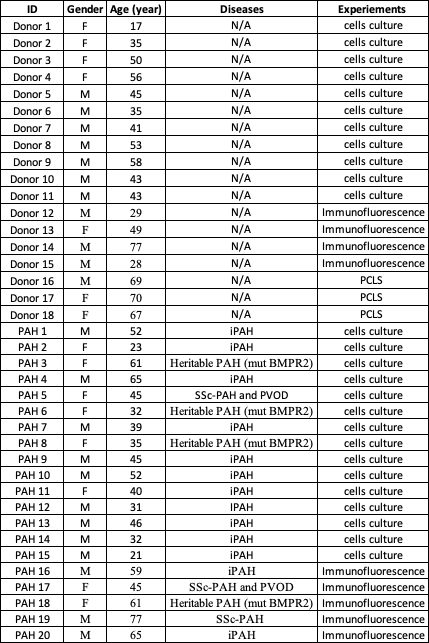

**Table S2. Characteristics of Donor and PAH patients**

*iPAH, idiopathic Pulmonary Artery Hypertension*

*Mut BMPR2, mutation BMPR2*

*SSC-PAH and PVOD, systemic Sclerosis-Associated Pulmonary Arterial Hypertension and Pulmonary veno-occlusive disease*

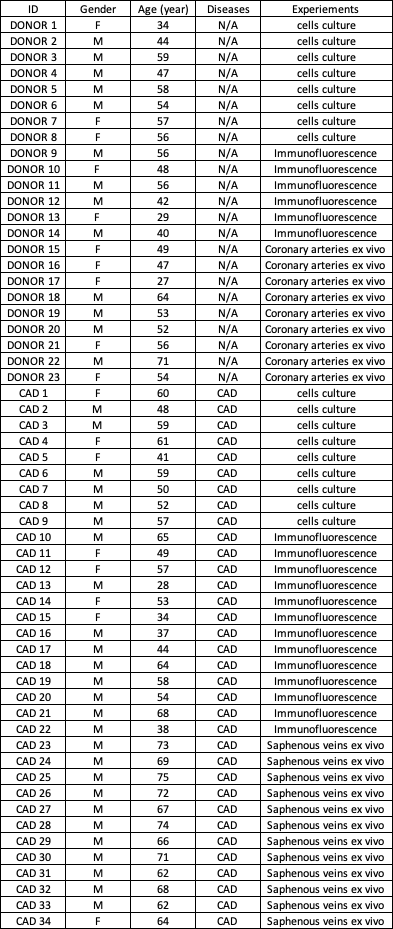

**Table S3. Characteristics of Donor and CAD patients**

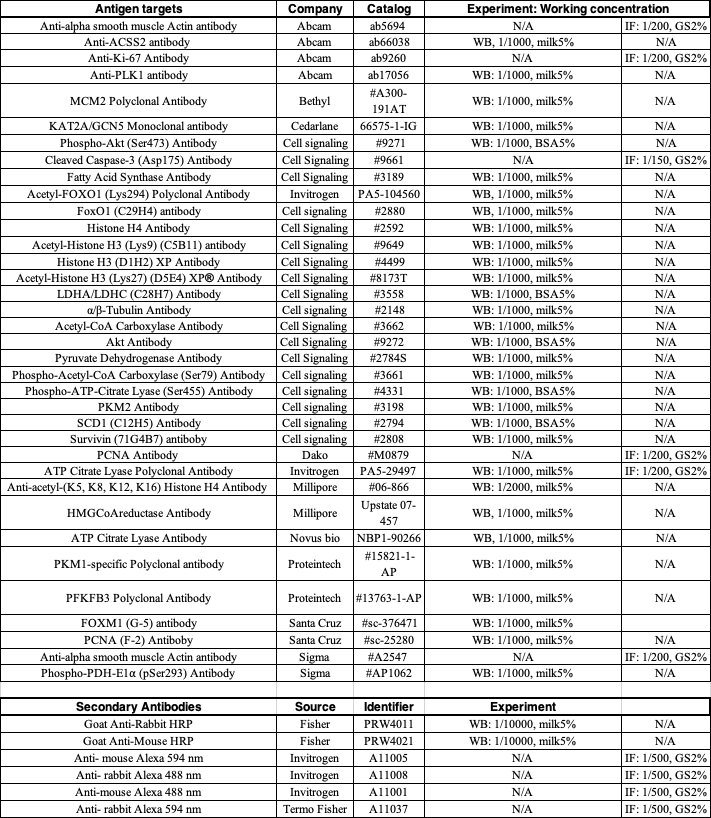

**Table S4. Antibody description**

*BSA, Bovine Serum Albumine*

*GS, Goat Serum*

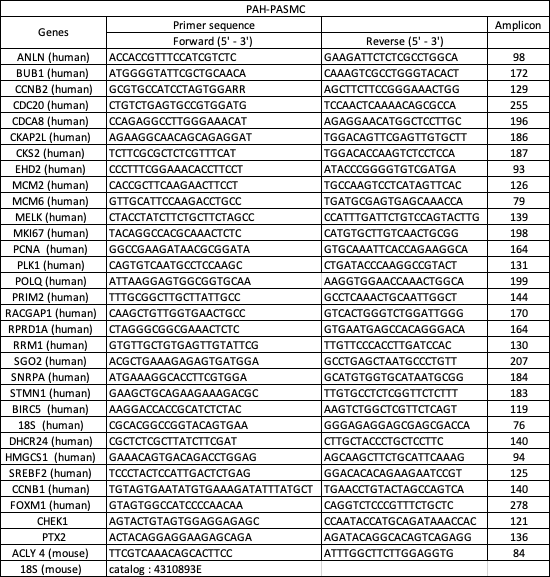

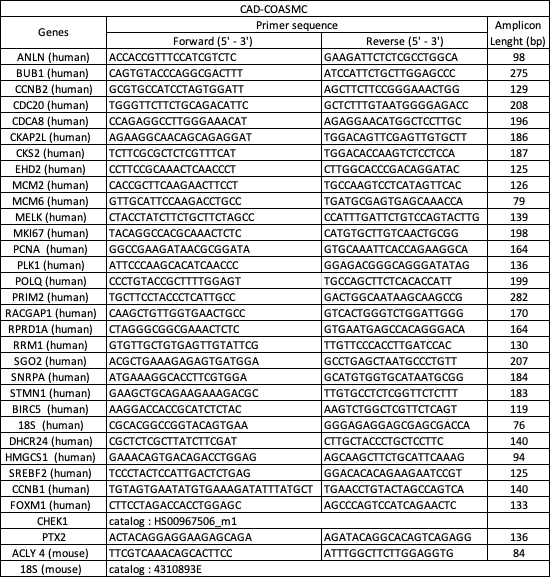

**Table S5. Primers description**
